# Differential Regulation of SIRT5 Activity by Reduced Nicotinic Acid Riboside (NARH)

**DOI:** 10.1101/2025.07.14.664716

**Authors:** Abu Hamza, Dickson Donu, Emily Boyle, Rasajna Madhusudhana, Alyson Curry, Yana Cen

## Abstract

SIRT5, one of the human sirtuins, catalyzes the removal of acyl substitutions from lysine residues in a NAD^+^-dependent manner. In addition to the deacetylase activity, SIRT5 also demonstrates strong desuccinylase, demalonylase, and deglutarylase activity. Through deacylating a broad spectrum of cellular proteins and enzymes, SIRT5 is heavily involved in the regulation of energy metabolism, reactive oxygen species (ROS) reduction, and ammonia detoxification. Accumulating evidence also suggest SIRT5 as a potential therapeutic target for the treatment of neurodegenerative diseases, metabolic disorders, and cancer. In the current study, we report the identification and characterization a SIRT5 modulator, reduced nicotinic acid riboside (NARH). It shows differential regulation of the distinct activities of SIRT5: activates desuccinylation, but mildly suppresses deacetylation. NARH binds to SIRT5 in the absence of NAD^+^, and demonstrates cellular target engagement and activity. The potential NARH binding site is further investigated using a suite of biochemical and computational approaches. The current study provides greatly-needed mechanistic understanding of SIRT5 regulation, as well as a novel chemical scaffold for further activator development.

## INTRODUCTION

Mammalian sirtuins are a group of “eraser” enzymes that remove acyl modifications from lysine residues in a NAD^+^-dependent fashion. The seven human sirtuins (SIRT1-SIRT7), with distinct subcellular localizations and biological functions, are the key players in various cellular events ranging from transcription regulation to energy homeostasis.^1^ The link between sirtuin activity and metabolism, first established by the discovery that these enzymes utilize NAD^+^,^2, 3^ has turned out to be more profound than originally envisioned. The ability of sirtuins to control metabolism by either modulating the transcription of or directly deacylating metabolic enzymes places sirtuins at the nexus of metabolic regulation.^4^

SIRT5, one of the mitochondrial sirtuins, harbors unique enzymatic activities compared with other members of the human sirtuin family. Although initially characterized as a deacetylase, SIRT5 possesses very weak or undetectable deacetylase activity *in vitro*.^5, 6^ Recent studies suggest that SIRT5 is a lysine desuccinylase, demalonylase and deglutarylase.^5, 7, 8^ It prefers to remove negatively charged acyl groups from lysine residues owing to the unique active site amino acid (AA) residues, Ala86, Tyr102 and Arg105.^5, 7^ Aided by viable SIRT5-deficient mouse models and mass spectrometry (MS)-based proteomic surveys, hundreds of SIRT5 cellular targets with succinylated, malonylated and/or glutarylated lysines have been identified.^8–11^ Despite these studies, the biological significance and regulation of the diverse SIRT5 activities remain largely elusive. For example, it has been proposed that acylation of many SIRT5 substrates occurs in a non-enzymatic fashion under the chemical conditions in the mitochondrial matrix, as in the case of lysine succinylation.^12^ Thus, the widespread protein lysine acylation in mitochondria may simply be the result of the high pH and acyl-CoA concentrations in this organelle.^12^ SIRT5 serves as the “quality control” enzyme to maintain mitochondrial health through removal of these acylation marks. Furthermore, although the breadth of SIRT5 targets is impressive, tissue-specific or conditional knockout of SIRT5 had marginal functional phenotypes except for the significant increase of lysine acylation which may have very little biological relevance.^13, 14^ Innovative chemical tools are needed to further deconvolute the biological functions of SIRT5.

There are only a handful of small-molecule SIRT5 activators discovered so far. Increased levels of NAD^+^, the co-substrate of sirtuin-catalyzed reactions, has been shown to upregulate SIRT5 activity.^15, 16^ Mammalian sirtuins including SIRT5 have relatively low binding affinity for NAD^+^ to ensure the transient elevation in NAD^+^ content produced by cellular events can be readily sensed. However, boosting NAD^+^ level is known to activate other human sirtuin isoforms as well.^17, 18^ Polyphenols such as resveratrol and piceatannol have been suggested to activate SIRT5 against artificial fluorophore-conjugated peptide substrates.^19^ But they fail to activate SIRT5 against its physiological substrates, and suffer from low solubility.^19^ Recently, MC3138 and its derivatives have emerged as SIRT5-selective activators.^20, 21^ Unfortunately, they are not particularly potent, and the low solubility again limits the complete characterization of these compounds. Our group has identified a water soluble and cell-permeable SIRT5 activator, nicotinamide riboside (NR).^22^ It selectively activates the deacetylase activity of SIRT5, but causes negligible changes to the desuccinylase activity.^22^

We here report the synthesis and characterization of another SIRT5 activator, reduced nicotinic acid riboside (NARH). Although initially generated as a potential NAD^+^ booster, NARH does not serve as a NAD^+^ precursor. Instead, it activates the desuccinylase activity of SIRT5 with either synthetic peptide or physiological substrates. Interestingly, the deacetylase activity of SIRT5 is rather inert to NARH, showing mild inhibition at high micromolar concentrations. The NARH-SIRT5 interaction was further investigated. NARH binds to SIRT5 directly with a binding affinity in the low micromolar range as determined by microscale thermophoresis (MST) and isothermal titration calorimetry (ITC). The cellular target engagement was confirmed by cellular thermal shift assay (CETSA). Mutagenesis studies revealed the potential NARH interacting amino acid residues (AAs). Our results illuminate key regulatory mechanisms of SIRT5 activity, and provide the blueprint for further activator development.

## RESULTS AND DISCUSSION

### Synthesis of NARH

Reduced nicotinamide riboside (NRH) is a newly discovered NAD^+^ precursor.^23^ A putative pathway has been proposed for the intracellular conversion of NRH to NAD^+^ (**Figure 1**).^24^ NRH can be phosphorylated by adenosine kinase (AK) to its corresponding mononucleotide, NMNH. Subsequently, the adenylylation of NMNH catalyzed by nicotinamide mononucleotide adenylyltransferase (NMNAT) leads to the formation of NADH which can be converted to NAD^+^ *via* redox reactions. NRH is an orally bioavailable and potent NAD^+^ precursor which holds great potential as a therapeutic agent for NAD^+^ restoration.^23, 25^

**Figure 1.**
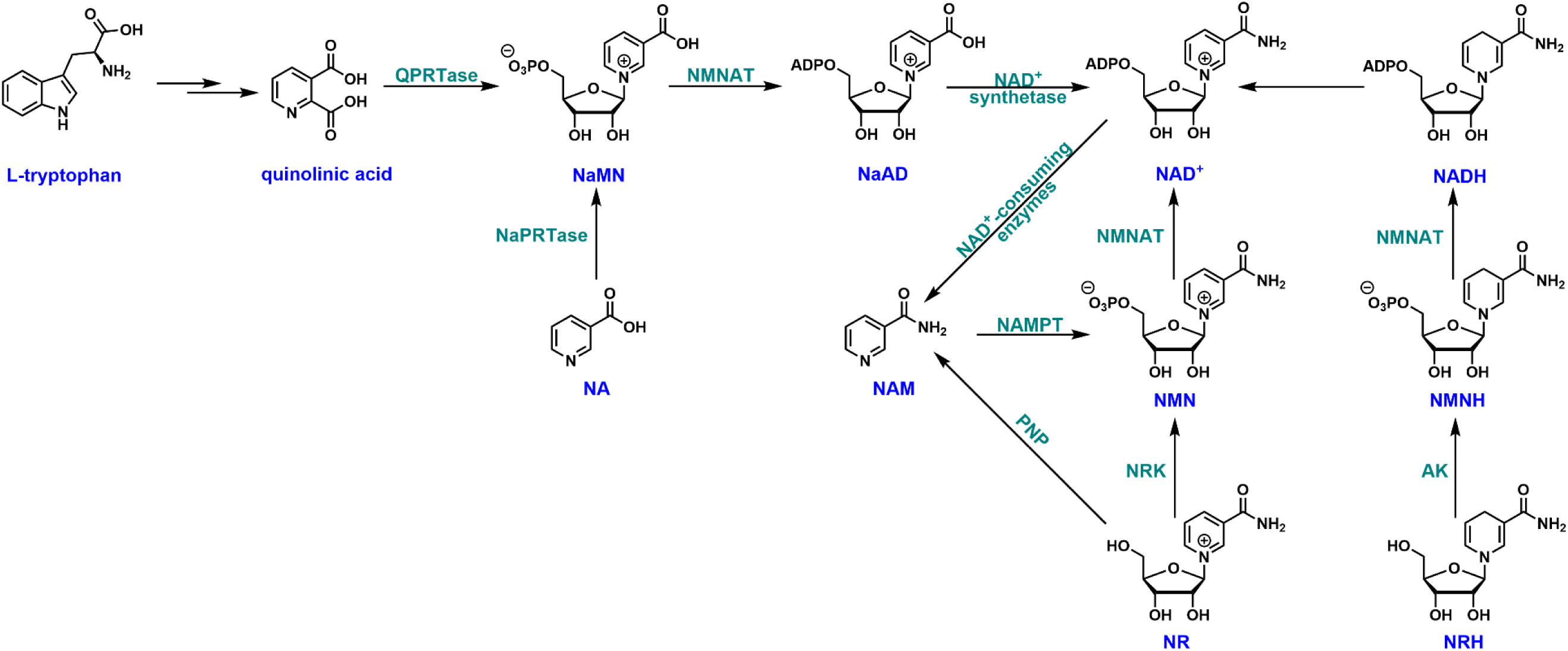
NAD^+^ biosynthetic pathways in mammalian cells.

Much attention has been devoted to the study of NRH, although the potential of NARH esters as NAD^+^ enhancement agents is enormous. We developed a convenient synthesis of the ethyl ester of NARH (**Figure 2**). Ethyl nicotinate was coupled with 1,2,3,5-tetra-*O*-acetyl-β-D-ribofuranose in the presence of TMSOTf.^26^ The “neighboring group participation” resulted in the formation of tri-*O*-acetylated β-ethylnicotinate riboside (intermediate A) in a stereoselective manner in quantitative yield. Subsequently, intermediate A was converted to the reduced intermediate B with Na_2_S_2_O_4_ in dichloromethane and saturated NaHCO_3_ mixture in 91% yield. Intermediate B was then unmasked in 4N ammonia in methanol at 4°C to afford the ethyl ester of NARH in 95% isolated yield. The entire synthetic sequence only required one column chromatography (the deprotection of intermediate B). It provided unprecedented easiness and high yield in generating NARH esters.

**Figure 2.**
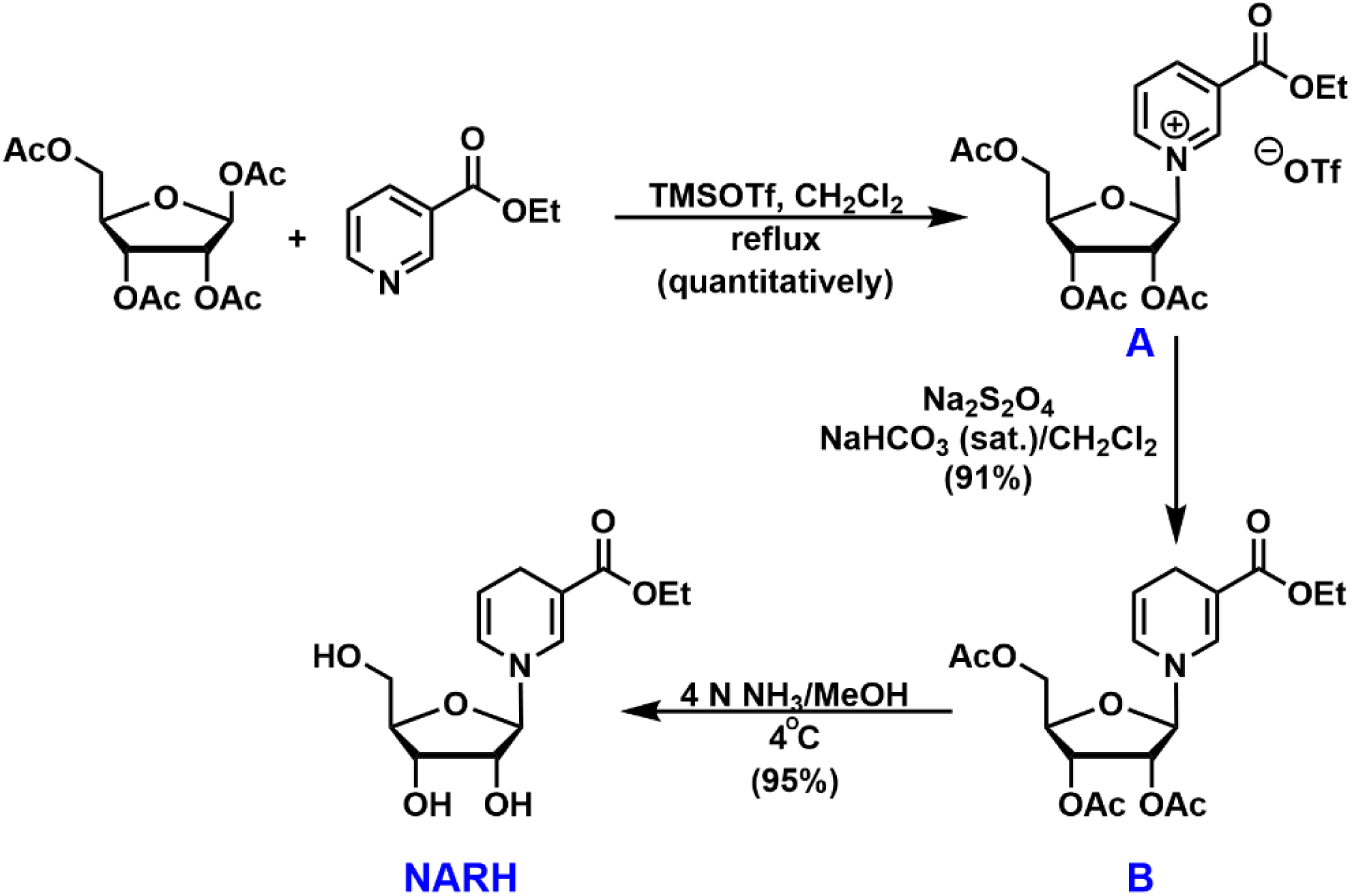
Synthesis of NARH.

### NARH activates SIRT5 desuccinylation

Numerous proteomic studies have elucidated a landscape of hundreds of SIRT5 substrates. However, these proteomic analyses also revealed that many of the succinylation sites were acetylated at the same position.^9,27^ With desuccinylation and deacetylation leading to the same unmodified lysine product, an intriguing question that deserves further investigation is: is there any intrinsic regulatory mechanism allowing SIRT5 to switch from one activity to another under different physiological conditions? Our first hint came from a study showing that SIRT5 activities demonstrated differential sensitivities to nicotinamide (NAM), a byproduct of sirtuin-catalyzed reactions.^28^ Physiological NAM concentrations potently inhibit the desuccinylase activity, but cause negligible changes on the deacetylase activity. In the same vein, we recently discovered that NR serves as SIRT5 activator.^22^ It stimulates SIRT5-mediated deacetylation of synthetic peptides or endogenous substrates, but fails to activate the desuccinylation to any appreciable levels. Encouraged by these prior findings, we further explored NARH as a potential SIRT5 regulator.

Our initial effort focused on the assessment of SIRT5 activity toward a synthetic peptide substrate derived from histone H3 (H3K9Suc) using an HPLC-based assay as described in “Methods and Materials”. Recombinant human SIRT5 demonstrated robust desuccinylase activity with a *K*_m_ value of 75.7 ± 3.8 μM, and a *k*_cat_ value of 0.029 ± 0.0014 s^-1^ (**Table 1**), consistent with previous reports.^5, 22^ NARH was able to increase the desuccinylation by almost 2.8-fold at 800 μM. Titration of NARH resulted in a concentration-dependent SIRT5 activation with an EC_50_ of 88 μM (**Figure S3**).

**Table 1.** Recombinant SIRT5 steady state parameters^a^.

| SIRT5 | Substrate | $K_m$ (μM) | $k_{cat}$ ($10^{-3}s^{-1}$) | M.A. (NARH)<sup>b,c</sup> |
| --- | --- | --- | --- | --- |
| **WT** | H3K9Suc | $75.7 \pm 3.8$ | $29.0 \pm 1.4$ | 2.8 |
| | p53K382Ac | $1058 \pm 109$ | $18.7 \pm 2.8$ | **0.73** |
| **Y102A** | H3K9Suc | $1230 \pm 170$ | $37.1 \pm 5.6$ | 1.1 |
| **R105A** | H3K9Suc | $616 \pm 100$ | $7.4 \pm 0.4$ | 0.95 |
| **W222A** | H3K9Suc | $193 \pm 41$ | $25.0 \pm 1.5$ | 1.5 |
| **A86S** | H3K9Suc | $139 \pm 26$ | $67.3 \pm 7.4$ | 0.41 |

<sup>a</sup> At saturating NAD<sup>+</sup> concentration. <sup>b</sup> [NARH] = 800 μM.
<sup>c</sup> Maximum Activation = $k_{cat}(act.) * K_m(unact.) / (k_{cat}(unact.) * K_m(act.))$

The isoform-selectivity of NARH was further evaluated. It was tested against recombinant SIRT1, SIRT2, SIRT3, and SIRT6 using the same HPLC assay and synthetic peptides as the substrates. No appreciable activity increase was detected for the other human sirtuin isoforms at up to 800 μM (**Figure S4**). The effect of NARH on physiological SIRT5 substrates was also evaluated. HeLa cell lysate was treated with recombinant SIRT5 and NAD^+^ in the presence or absence of NARH. The succinylation levels of carbamoyl phosphate synthase 1 (CPS1), the rate-limiting enzyme in the urea cycle and a known endogenous SIRT5 substrate,^5, 15^ was probed by western blot. As shown in **Figure 3A**, NARH treatment at 800 μM led to a reduction of the succinylation level of CPS1, presumably due to SIRT5 activation. Furthermore, the mitochondrial lysate from HEK293 cells was incubated with SIRT5 with or without NARH. In the presence of NARH, the succinylation level of multiple protein bands showed significant decrease as compared to the no NARH controls (**Figures 3B** and **3C**), suggesting the activation of SIRT5 desuccinylase activity by NARH on physiological substrates.

**Figure 3.**
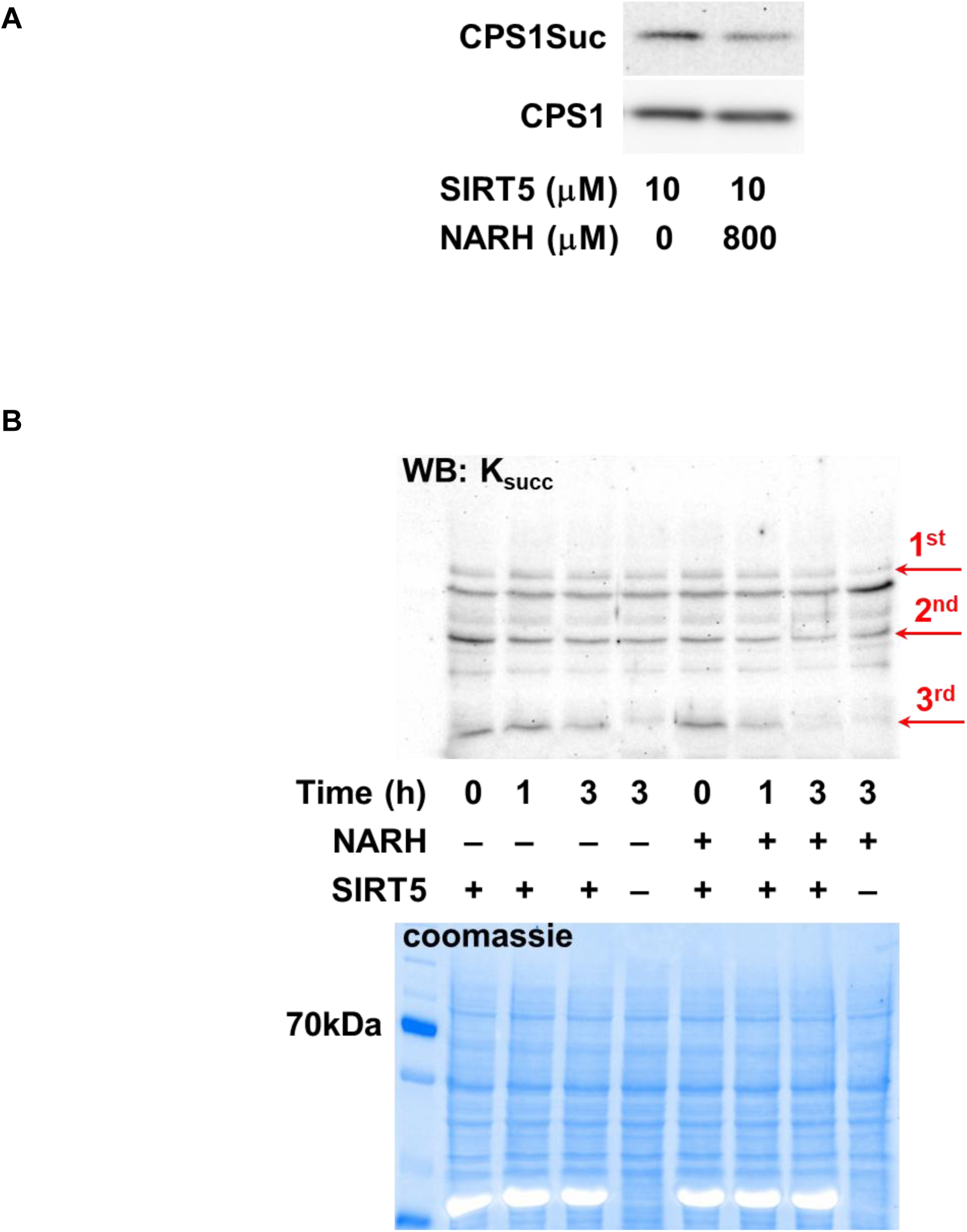
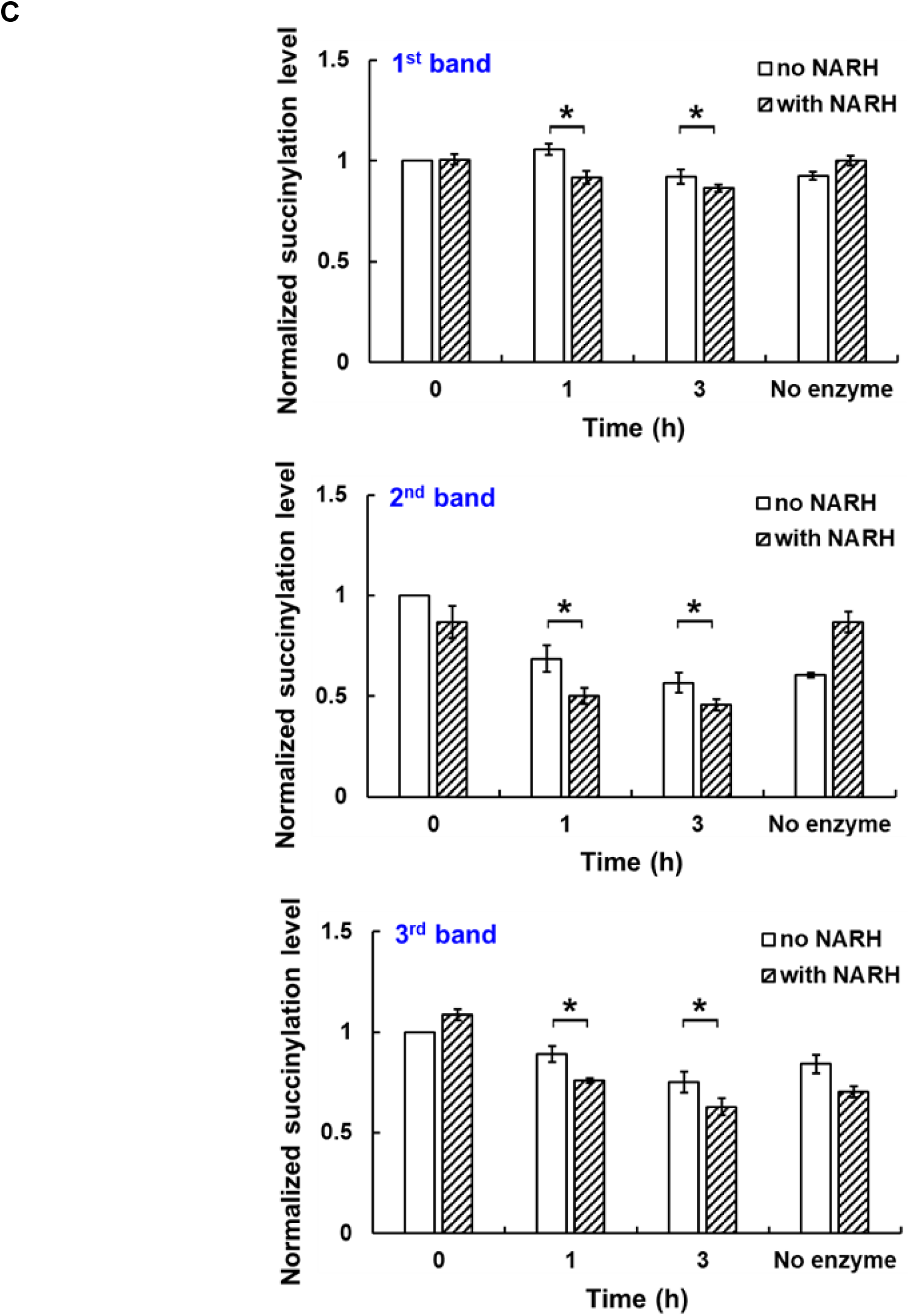
Activation of SIRT5 by NARH on physiological substrates. (A) Western blot showing decreased succinylation level of CPS1 in HeLa cell lysate treated with NARH; (B) Representative western blot (top) showing the desuccinylase activity of SIRT5 on HEK293 cell mitochondrial lysate in the presence or absence of NARH. Coomassie staining (bottom) serves as the loading control; (C) Quantification of the western blot results in (B). The relative lysine succinylation level was calculated by setting the level of no NARH control zero time to 100%. The quantification data represents the average of three independent experiments ± SD. Statistical significance was determined by a Student’s *t*-test: \**p* < 0.05 *vs* no NARH control.

### NARH binds to SIRT5 directly

NARH binding to SIRT5 in the absence of any substrate was analyzed by microscale thermophoresis (MST) experiments to reveal a *K*_d_ value of 6.1 ± 0.5 μM (**Figure 4A**). To better understand the direct binding of NARH to SIRT5, the dissociation constant was also measured by isothermal titration calorimetry (ITC). NARH bound to SIRT5 with a *K*_d_ of 8.3 ± 2.8 μM (**Figure 4B**), in good agreement with the MST results. Interestingly, NARH bound with a negative -*TΔS*, suggesting that there is more disorder once NARH binds. Consistent with a previous report, SIRT5 did not bind to NAD^+^ in the absence of the peptide substrate (**Figure S5A**).^29^ Furthermore, in the presence of NAD^+^, NARH bound to SIRT5 with a *K*_d_ of 6.6 ± 1.5 μM (**Figure S5B**), similar to that in the absence of NAD^+^. All these results suggested that NARH may bind to a distinct allosteric site to stimulate desuccinylation *via* either a conformational change or an influence of the enzyme dynamics.

**Figure 4.**
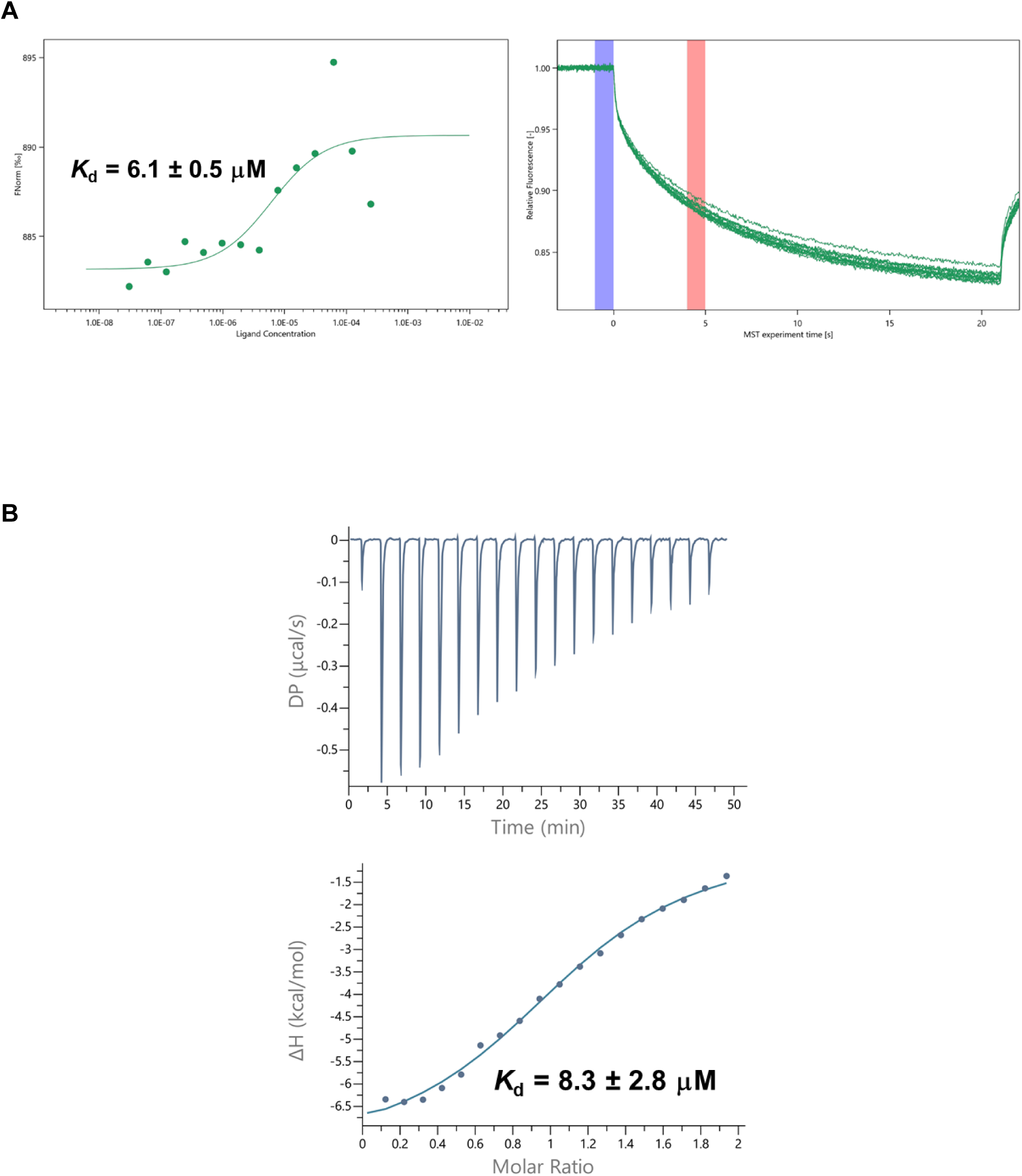
NARH binds to SIRT5 in the absence of NAD^+^. (A) Determination of binding affinity of NARH to SIRT5 using MST. Left: raw MST traces; Right: binding isotherms for MST signals *vs* NARH concentration; (B) Representative ITC trace of NARH titrated into SIRT5. The *K*_d_ was determined to be 8.3 ± 2.8 μM.

### Regulation of SIRT5 desuccinylase/demalonylase activity by NARH in cells

Small molecule modulators can serve both as useful tools for interrogating protein biological functions and as therapeutic agents. However, maintaining the desirable on-target effect of a small molecule in a cellular context can be challenging. The target engagement of NARH was evaluated by cellular thermal shift assay (CETSA). CETSA conditions were first optimized using recombinant SIRT5. As shown in **Figure 5A**, the abundance of soluble SIRT5 decreased with increasing temperature, showing that the thermostability of SIRT5 can be monitored by this method. Importantly, upon incubation with NARH, the stability of SIRT5 was increased with a Δ*T*_m_ of 3.3 °C (**Figure 5B**), indicating the physical interaction of NARH with SIRT5.

**Figure 5.**
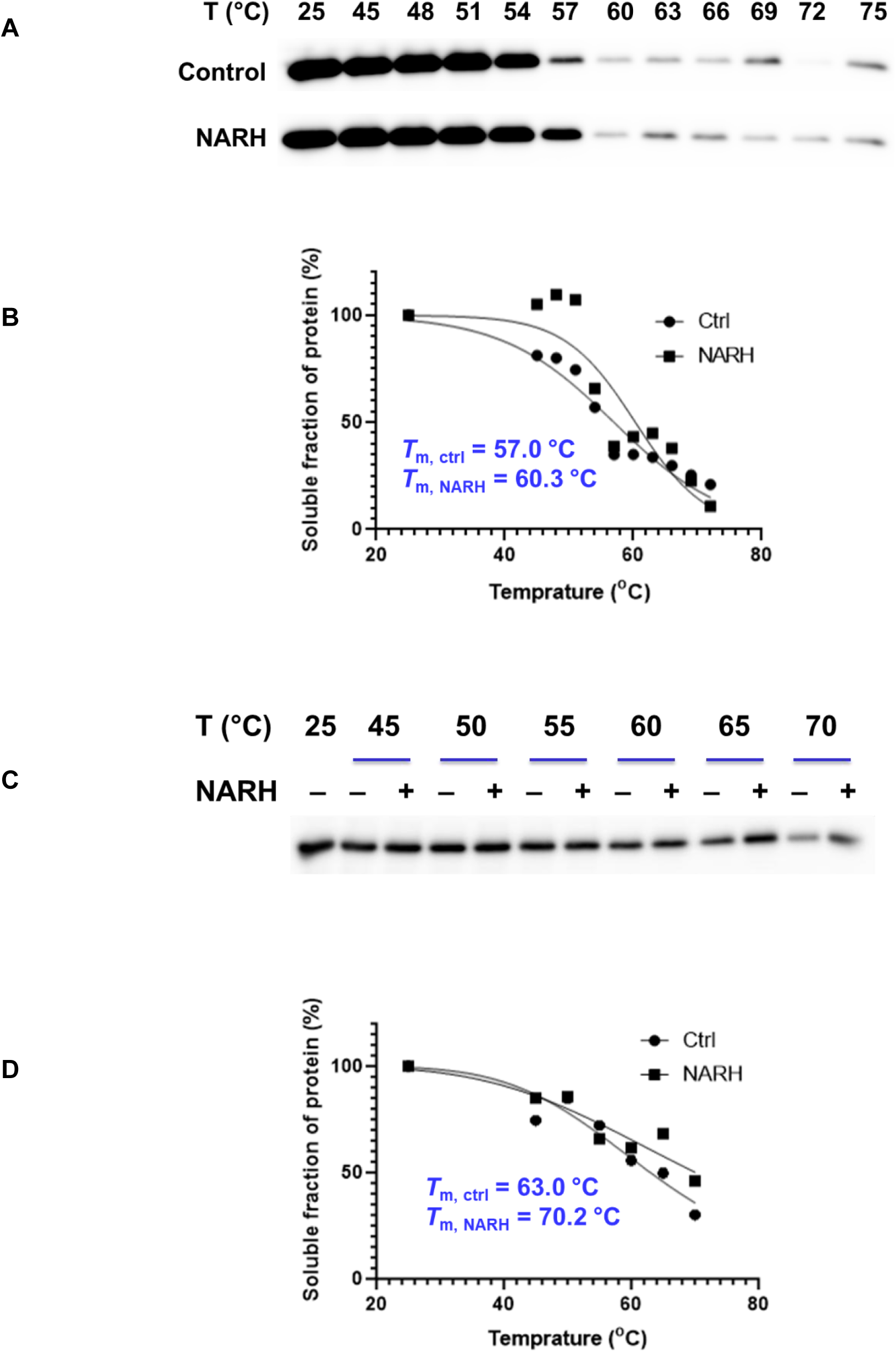
Target engagement of NARH assessed by CETSA. (A) Representative western blots of thermal shift assay using recombinant SIRT5; (B) melt curves for control and NARH-treated recombinant SIRT5; (C) Representative western blots of thermal shift assay using SIRT5-overexpressing cell lysate; (D) CETSA curves for control and NARH-treated cell lysates.

Subsequently, the lysate of HEK293 cells overexpressing Flag-SIRT5 was incubated in the presence or absence of NARH, and heated as described in “Methods and Materials”. After centrifugation, the soluble fractions were resolved by SDS-PAGE and analyzed by immunoblot. The thermal stability of SIRT5 was enhanced upon NARH treatment with a Δ*T*_m_ of 7.2 °C (**Figures 5C** and **5D**). These results confirmed that SIRT5 is a direct binding target of NARH in cell lysate. To investigate the cellular activity of NARH, a metabolic labeling strategy using a malonate derivative, MalAM-yne^30^ (**Figure 6A**), was employed. Previous studies have shown that this compound can be incorporated into cellular proteins, primarily through lysine malonylation, and that MalAM-yne–modified proteins serve as substrates for SIRT5.^30^ HeLa cells were pre-treated with either vehicle or 1 mM NARH, followed by incubation with 200 μM MalAM-yne (**Figure 6B**). The labeled proteins can be visualized by Cu (I)-mediated “click” conjugation with TAMRA-azide. As shown in **Figure 6C**, NARH treatment significantly enhances the removal of MalAM-yne from labeled proteins, supporting the notion that NARH functions as a SIRT5 activator.

**Figure 6.**
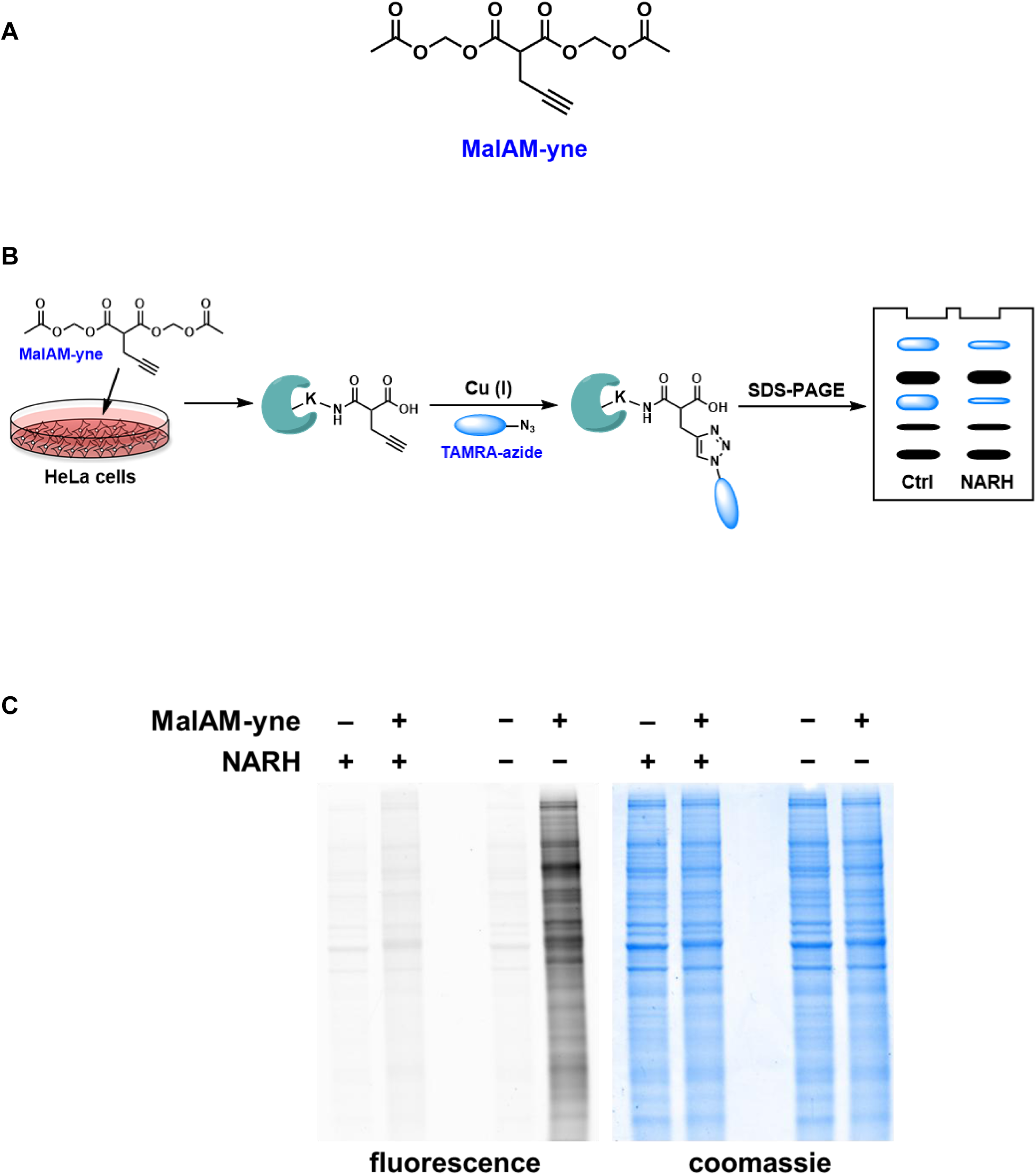
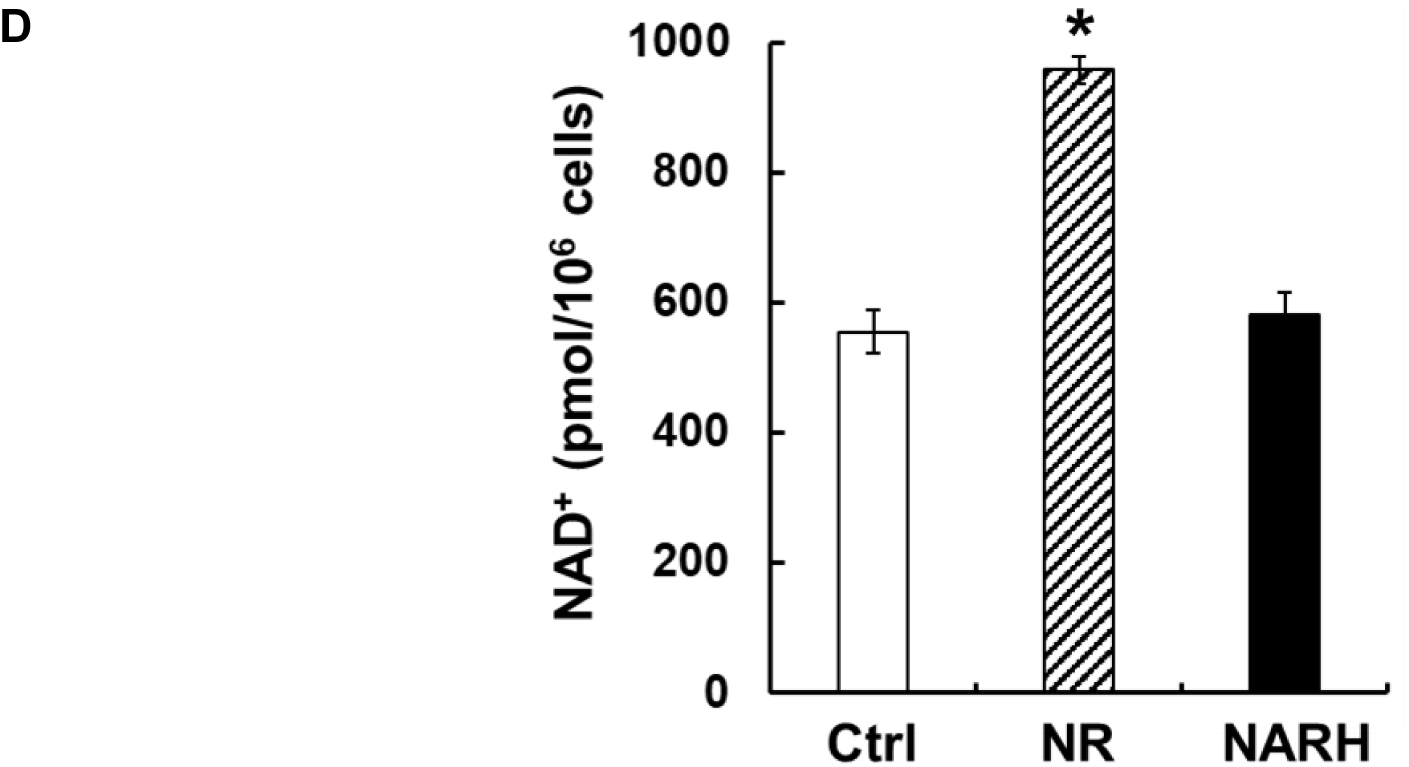
Activation of SIRT5 by NARH in cells. (A) Chemical structure of MalAM-yne; (B) Schematic representation of the metabolic labeling; (C) Analysis of malonylation using MalAM-yne. HeLa cells were cultured with or without 1 mM NARH, followed by incubation in the presence or absence of 200 μM MalAM-yne. The labeled proteins were “click” conjugated to TAMRA-azide. Pre-incubation with NARH led to significantly reduced labeling compared to no NARH control; (D) Intracellular NAD^+^ levels in control, NR (1 mM), or NARH (1 mM)-treated HEK293 cells. The quantification data represents the average of three independent experiments ± SD. Statistical significance was determined by a Student’s *t*-test: \**p* < 0.05 *vs* control.

NARH was initially synthesized as a potential NAD^+^ boosting agent. Elevated cellular NAD^+^ concentration has been shown to stimulate SIRT5 activity.^15^ It thus became critical to clarify whether the observed SIRT5 activity upregulation was due to direct activation by NARH or indirect activation *via* NAD^+^ increase. Incubation of HEK293 cells with 1 mM nicotinamide riboside (NR), a known NAD^+^ precursor, led to a nearly two-fold increase of intracellular NAD^+^ content (**Figure 6D**), consistent with previous reports.^16, 26^ Interesting, culturing the cells with up to 1 mM NARH caused negligible changes to the intracellular NAD^+^ levels (**Figure 6D**). Similar results were obtained in Neuro2a and HeLa cells (**Figure S6**). A recent study demonstrated that NARH treatment alone resulted in insignificant changes to NAD^+^ contents in INS1E and MEF cells as well as kidney and muscle,^31^ in good agreement with our data. Overall, unlike its structural analog NR, NARH does not serve as a NAD^+^ precursor under the conditions we tested, confirming the direct activation of SIRT5 by NARH in the cellular setting.

### SIRT5 deacetylation is insensitive to NARH

SIRT5 harbors unique NAD^+^-dependent deacetylase and deacylase activities.^32^ It has been implicated in the regulation of various metabolic pathways by removing acyl modifications from a broad array of endogenous targets.^8–11^ The promiscuity of SIRT5 has led to the hypothesis that the distinct activities of this enzyme can be differentially regulated by small molecules. The *k*_cat_ and *K*_m_ values of SIRT5-catalyzed deacetylation were determined with a synthetic peptide (**Table 1**), p53K382Ac. Indeed, the enzyme exhibited a rather weak catalytic efficiency with a *k*_cat_/*K*_m_ of 17.7 M^-1^s^-1^. Interestingly, the addition of 800 μM NARH only caused mild inhibition of the deacetylase activity with a *k*_cat_/*K*_m_ of 12.9 M^-1^s^-1^ (**Table 1**).

Differential regulation of SIRT5 activities has been previously reported. The desuccinylase activity is highly sensitive to NAM inhibition, whereas the deacetylase activity is NAM resistant.^28^ Our recent study indicated that NR preferentially activates the deacetylase activity, but not the desuccinylase activity.^22^ The current study adds a new dimension by revealing a distinct sensitivity pattern to NARH. These studies suggest that that SIRT5 “senses” the availability of small molecules and selectively engage its enzymatic activities in response to these molecular cues.

### Characterization of NARH and SIRT5 interactions

In order to better understand this unique regulatory mechanism, the interactions between NARH and SIRT5 were further characterized. In our previous NR report, a putative NR binding site involving Y102, R105 and W222 has been proposed based on molecular docking analysis and mutagenesis studies.^22^ Due to the structural similarity between NR and NARH, the same binding site was hypothesized as the potential NARH binding site. The steady-state kinetic parameters of Y102A, R105A and W222A mutants are shown in **Table 1**. Both Y102A and R105A exhibited significantly reduced desuccinylase activity as compared to the wtSIRT5, stemming primarily from increased *K*_m_ values. The R105A mutant also suffered from markedly decreased *k*_cat_, only one-fourth of the wildtype value. The results were consistent with the critical role these two AAs play in the preferred desuccinylation activity.^5^ The W222A mutant showed a nearly 3-fold decrease in catalytic efficiency in comparison with wtSIRT5, owing mainly to a 2.5-fold increase of *K*_m_ (193 ± 41 μM).

The binding affinity of the mutants with NARH was then evaluated by MST. Y102A demonstrated a complete loss of NARH binding (**Figure S7A**), while R105A had a relatively weak binding affinity (*K*_d_ = 130.2 ± 14 μM, **Figure S7B**) compared to the wildtype (*K*_d_ = 6.1 ± 0.5 μM). Furthermore, at 800 μM, NARH failed to activate the desuccinylase activity of the Y102A and R105A mutants to any appreciable levels (**Table 1**), thus confirming that Y102 and R105 are essential for the recognition and binding of NARH. The W222A mutant, on the other hand, bound to NARH with a higher affinity (*K*_d_ = 1.2 ± 0.3 μM, **Figure S7C**). The desuccinylation of W222A was rather inert to NARH, showing merely 1.5-fold activation at 800 μM concentration (**Table 1**). We reasoned that the Trp to Ala mutation reduced steric hindrance for an improved NARH binding. However, the same mutation may also cause loss of important hydrophobic interactions, leading to the insensitivity to NARH.

Another distinct active site AA for SIRT5 is A86. It is crucial for substrate interaction and is unique for SIRT5 because it is replaced by a Phe in other human sirtuin isoforms.^33, 34^ The presence of the sterically less hindered A86 allows SIRT5 to better accommodate large acyl modifications such as succinyl or malonyl groups. The A86S mutant still exhibited a strong desuccinylase activity with a *K*_m_ of 139 ± 26 μM and a *k*_cat_ of 0.067 ± 0.0074 s^-1^, comparable to the wtSIRT5. Strikingly, NARH acted as an inhibitor for A86S, causing a 2.5-fold inhibition at 800 μM (**Table 1**). The type of inhibition was further characterized. Lineweaver-Burk plot showed that NARH was noncompetitive with succinylated peptide substrate as evidenced by a series of lines that intersect at the x-axis (**Figure 7A**). Moreover, NARH demonstrated direct binding to A86S with a *K*_d_ value of 21.3 ± 3.5 μM as determined by MST (**Figure 7B**).

**Figure 7.**
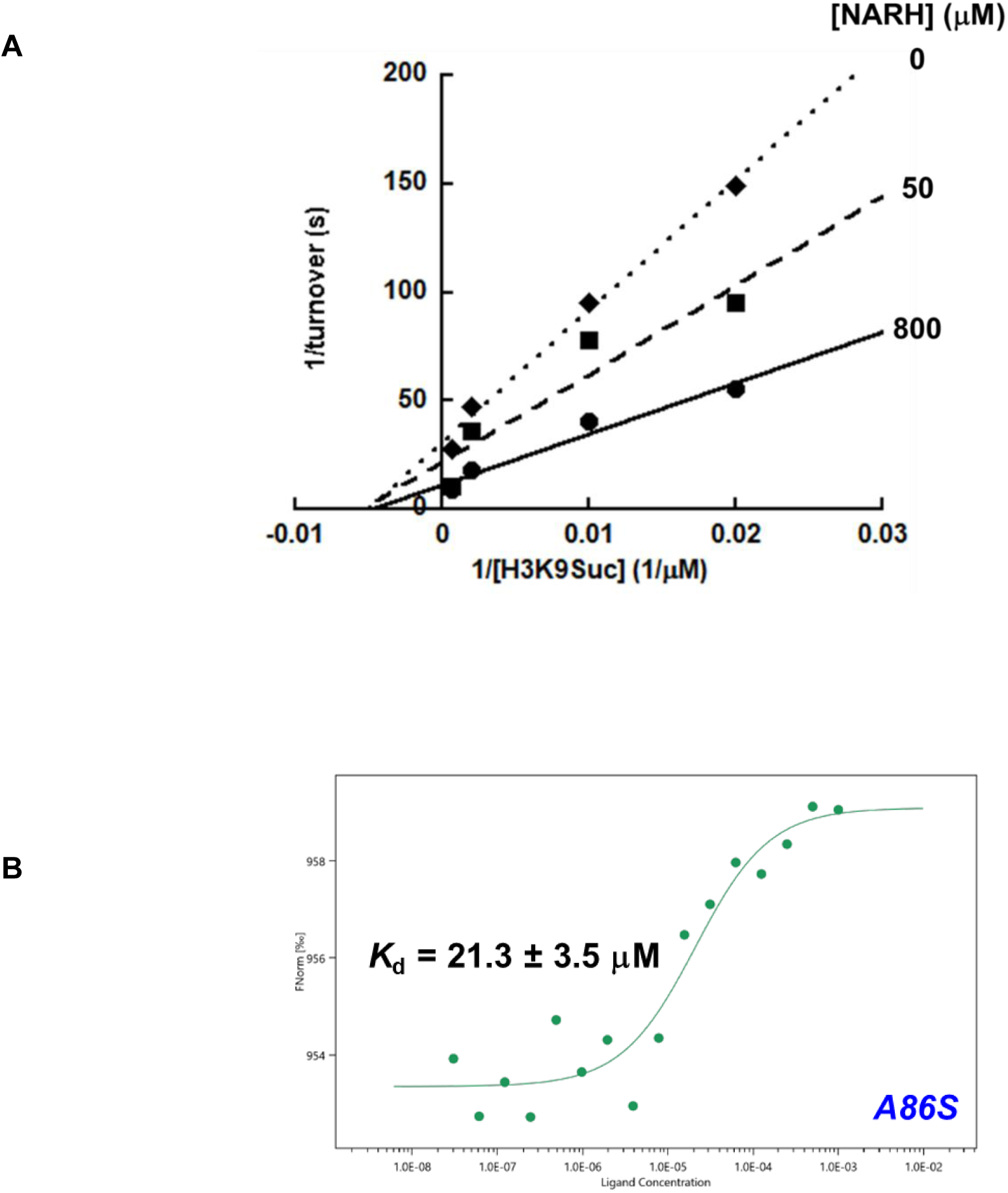
NARH is an inhibitor of the desuccinylase activity of A86S mutant. (A) Double-reciprocal plot showing NARH is a noncompetitive inhibitor of A86S; (B) Determination of binding affinity of NARH to A86S using MST.

Molecular docking analysis was also performed to further characterize the interactions of NARH with SIRT5 using the crystal structure of SIRT5 in complex with NAD^+^ and H3K9Suc peptide (PDB: 3RIY). NARH was docked into the binding site with GOLD^35^ and scored with the built-in ChemPLP scoring function. The highest scored pose was chosen as the optimal docking pose for further analysis. As shown in **Figure 8A**, NARH occupies a site in the cleft between the small zinc-binding domain and the large Rossmann fold domain, exposing the ethyl ester moiety to solvent. The molecule is sandwiched between one of the helices of the zinc-binding domain and the flexible loop^36^ connecting the two domains. The dihydropyridine ring of NARH forms hydrophobic interactions with the aromatic side chains of F101 and Y104 (**Figure 8B**). In addition, R105 makes H-bond interactions with the 3’- and 5’-OH groups. Y104 makes H-bond interaction with the ester carbonyl oxygen. Also notable is that the side chain of P190 forms hydrophobic interaction with the ethyl group of NARH.

**Figure 8.**
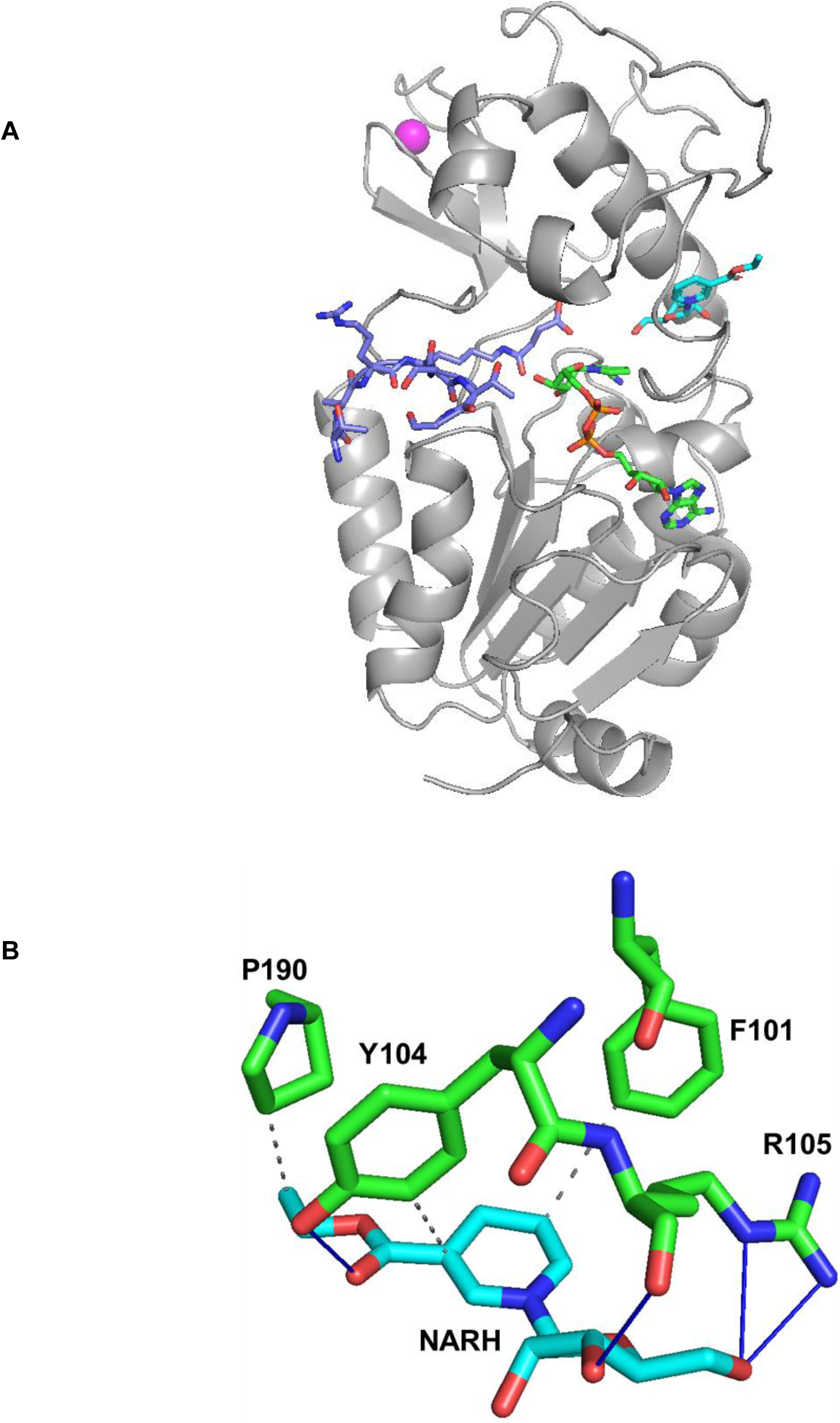
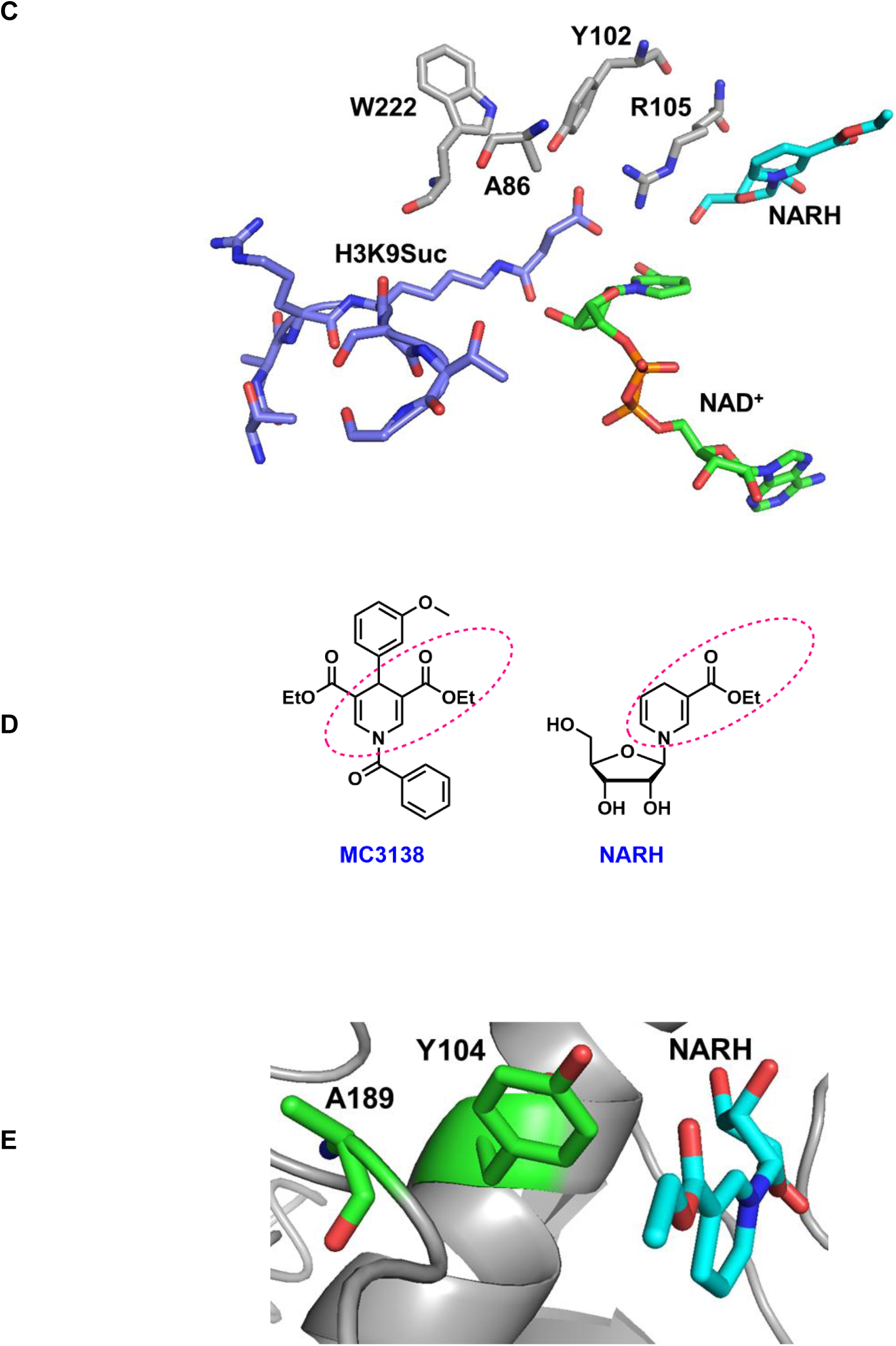
Docking analysis of NARH-SIRT5 interactions. (A) Docked pose of NARH in the allosteric binding site. NAD^+^: green; NARH: cyan; H3K9Suc: purple; zinc: magenta; (B) Interactions of NARH with amino acids in the binding site. H-bond: blue line; hydrophobic interaction: grey dashed line; (C) Overview of NARH, NAD^+^, and H3K9Suc in the docked pose. AAs: light grey; (D) Chemical structures of MC3138 and NARH. The common structural motif is highlighted in the pink dotted circles; (E) Relative position of NARH, Y104, and A189 in the docked pose.

The docked pose suggested that NARH occupies an allosteric site that is distal from the H3K9Suc binding site (**Figure 8C**). It is reasonable to speculate that the binding of NARH at this site induces conformational change in the substrate binding pocket for improved enzymatic turnovers. In the case of the A86S mutant, upon binding to NARH, the structural rearrangement may cause the sterically bulkier serine to clash with the succinyl group, consistent with the noncompetitive inhibition observed in the kinetic study.

MC3138 is a recently discovered SIRT5 activator.^20^ MC3138 treatment in cells mimic the effect of SIRT5 overexpression. There is certain structural similarity between MC3138 and NARH: both feature an ethyl-1,4-dihydropyridine-3-carboxylate moiety (**Figure 8D**). Moreover, docking study has placed MC3138 in a site near Y104 and A189.^20^ In our docked pose, NARH is situated at a similar site (**Figure 8E**), further suggesting that both compounds may share the same mechanism of action. Currently, we are applying structural biology approaches to reveal the NARH binding site, and to provide basis for future activator development.

## CONCLUSIONS AND PERSPECTIVES

SIRT5, one of the human sirtuins, has attracted significant attention during the last few years, primarily due to its unique enzymatic functions. Initially classified as a protein lysine deacetylase, SIRT5 demonstrates poor catalytic efficiency against acetylated peptide substrates, several orders of magnitude lower than the those of SIRT1, SIRT2, and SIRT3.^5^ This has led to the hypothesis that maybe SIRT5 has evolved away from being a deacetylase, maybe it has other activities. Indeed, later studies suggested that SIRT5 is a lysine desuccinylase, demalonylase, and deglutarylase.^5, 7, 8^ It prefers to remove acyl groups with negative charges. The discovery of these novel activities has greatly expanded the substrate repertoire of SIRT5, and deepened our understanding of the biological functions of this enzyme. The plethora of activities give SIRT5 its far-reaching role in maintaining metabolism and nutrient homeostasis.^32^ Additionally, SIRT5 has been implicated in the regulation of various pathologies such as cancer^20, 37^ and neurodegeneration.^38^ There is a clear need for small molecule SIRT5 regulators for both the functional annotation and pharmacological perturbation.

In the current study, we identified a SIRT5-selective activator, NARH. It is a structural analog of NRH, a known NAD^+^ precursor.^23^ However, unlike NRH, NARH treatment does not increase cellular NAD^+^ contents to any appreciable levels. Rather, it stimulates the desuccinylase activity of SIRT5 with both synthetic peptide and endogenous substrates. The deacetylase activity of SIRT5, on the contrary, is insensitive to NARH. This activation seems to be very specific for SIRT5 as NARH fails to activate several other human sirtuins. Biophysical analysis indicates that NARH binds to SIRT5 directly with a *K*_d_ value in the low micromolar range. Furthermore, NARH demonstrates cellular activity to decrease the succinylation levels of CPS1 in HeLa cells, and exhibits target engagement as shown by CETSA. The potential NARH binding site has been explored using a combination of site-directed mutagenesis, enzyme kinetics analysis, and docking studies. It has been proposed to locate in the cleft between the zinc-binding domain and Rossmann fold domain, near the NAD^+^ binding loop.^33, 36^ This putative allosteric site can be further exploited for the development of more potent and selective SIRT5 activators.

Overall, NARH is an isoform-selective SIRT5 activator with good druglike properties as predicted by SwissADME (**Figure S8**).^39^ The current study not only provides a lead compound for development of activators with improved potency and efficacy, but also generates novel mechanistic understanding on how allosteric site of SIRT5 can be targeted for activity regulation.

## METHODS AND MATERIALS

### Reagents and Instruments

All reagents were purchased from Aldrich or Fisher Scientific and were of the highest purity commercially available. HPLC was performed on a Dionex Ultimate 3000 HPLC system equipped with a diode array detector using Macherey-Nagel C18 reverse-phase column. NMR spectra were acquired on a Bruker AVANCE III 500 MHz high-field NMR spectrometer and the data were processed using Topspin software. HRMS spectra were acquired with either a Waters Micromass Q-tof Ultima or a Thermo Scientific Q-Exactive hybrid Quadrupole Orbitrap.

### Synthetic Peptides

Synthetic peptides H3K9Ac: ARTKQTAR(K-Ac)STGGKAPRKQLAS, p53K382Ac: KKGQSTSRHK(K-Ac)LMFKTEG, and H3K9Suc: ARTKQTAR(K-Suc)STGGKAPRKQLAS were synthesized and purified by Genscript. The peptides were purified by HPLC to a purity >95%.

### Protein Expression and Purification

Plasmids of SIRT1 (full length), SIRT2 (38-356), SIRT3 (102-399), SIRT5 (34-302), and SIRT6 (1-314), were generous gifts from Dr. Hening Lin (Cornell University). The proteins were expressed and purified according to previously published protocols.^40^ NRK1 was a gift from Cheryl Arrowsmith (Addgene plasmid # 60109). The protein was expressed and purified as described before.^41^ The identity of the protein was confirmed by tryptic digestion followed by LC-MS/MS analysis performed at the Vermont Biomedical Research Network (VBRN) Proteomics Facility. Protein concentrations were determined by Bradford assay.

### Deacetylation/Desuccinylation Assay

The *K*_m_ and *k*_cat_ of SIRT5 were measured for synthetic peptide substrates. A typical reaction was performed in 100 mM phosphate buffer pH 7.5 in a total volume of 50 μL. The reactions contained various concentrations of peptide substrate and 800 μM NAD^+^. Reactions were initiated by the addition of 10 μM of SIRT5 and were incubated at 37 °C for 20 min (deacetylation) or 5 min (desuccinylation) before being quenched by 8 μL of 10% TFA. The samples were then injected on an HPLC fitted to a Macherey-Nagel Nucleosil C18 column. Acylated and deacylated peptides were resolved using a gradient of 10%–40% acetonitrile in 0.1% TFA. Chromatograms were analyzed at 215 nm. Reactions were quantified by integrating area of peaks corresponding to acylated and deacylated peptides. Rates were plotted as a function of substrate concentration and best fits of points to the Michaelis-Menten equation were performed by Kaleidagraph®.

### Lineweaver-Burk Double-reciprocal Plot Analysis

Peptide titration reactions containing either 0, 50 μM, or 800 μM NARH were incubated with 800 μM NAD^+^, varying concentrations of H3K9Suc in 100 mM phosphate buffer pH 7.5. Reactions were initiated by the addition of 10 μM of SIRT5 and were incubated at 37 °C for 10 min before being quenched by 8 μL of 10% TFA. The samples were then injected on an HPLC fitted to a Macherey-Nagel Nucleosil C18 column. Acylated and deacylated peptides were resolved using a gradient of 10%–40% acetonitrile in 0.1% TFA. Chromatograms were analyzed at 215 nm. Reactions were quantified by integrating area of peaks corresponding to acylated and deacylated peptides. Double reciprocal plots were generated using Kaleidagraph® and fit to a linear curve representative of the Lineweaver-Burk relationship.

### Microscale Thermophoresis (MST) Binding Assay

The MST experiments were performed using a Monolith NT.115 instrument (NanoTemper Technologies). The purified recombinant His-tagged SIRT5 was labeled by the RED-tris-NTA 2^nd^ generation dye (NanoTemper Technologies). The protein concentration was adjusted to 200 nM in PBS-T buffer, while the dye concentration was set to 100 nM. Equal volumes (100 μL) of protein and dye solutions were mixed and incubated at room temperature in the dark for 30 min. Sixteen solutions of NARH with decreasing concentrations were prepared in the MST buffer by serial half-log dilution. Each solution was mixed with an equal volume of the labeled SIRT5. The binding affinity analysis was performed using standard MST capillaries, 40% excitation power (Nano-RED) and medium MST power at room temperature. The *K*_d_ value was determined with the MO.Affinity Analysis software (NanoTemper Technologies), using three independent MST measurements.

### Isothermal Titration Calorimetry

Experiments were performed at 25 °C using a MicroCal iTC200 titration calorimeter (Malvern Panalytical). The wtSIRT5 and mutant proteins were dialyzed into ITC buffer containing 20 mM Tris-HCl, pH 8.0, 100 mM NaCl, 100 μM TCEP, and 5% glycerol for 48 h before experiments were conducted. The titration experiments were performed as follows: 50 μM SIRT5 was injected into the sample cell using a loading syringe, with a solution of 500 μM NARH loaded into the injection syringe. The sample cell received one preliminary injection of 0.5 μL of the ligand followed by 18 injections of 2 μL ligand. Binding data for all experiments were analyzed using the ORIGIN 7.0 software (OriginLab) using a single set of binding sites to calculate the binding affinity, stoichiometry (N), and the thermodynamic values. *Cell Culture* HEK293 and HeLa cells were cultured in DMEM supplemented with 10% fetal bovine serum (FBS), 100 U/mL penicillin and 100 mg/mL streptomycin. Cells were maintained in a humidified 37°C incubator with 5% CO_2_.

### Cell Lysate Preparation

Cells were harvested and lysed with RIPA buffer (Thermo Fisher Scientific) supplemented with protease inhibitor cocktail (Thermo Fisher Scientific). Protein concentration was determined by Bradford assay.

### Cellular Thermal Shift Assay (CETSA)

The cell lysate was incubated with buffer or NARH for 30 min at room temperature. Subsequently, the samples were heated for 3 min at ten different temperatures with endpoints spanning 45°C∼72°C. Immediately after heating, the samples were incubated at room temperature for 3 min and then centrifuged to pellet the precipitated and aggregated proteins. The supernatant was transferred to a new tube. The soluble fraction was further analyzed by western blot.

### Immunoprecipitation

Cell lysate was incubated with anti-CPS1 antibody (Abcam) overnight at 4 °C. Then, protein A sepharose beads were added and incubated for 1 h. The beads were rinsed, and boiled with sample buffer for the subsequent western blotting.

### Western Blot

The cell lysate was resolved on a 10% SDS-PAGE gel and transferred to Immobilon PVDF transfer membrane (Biorad). The blot was blocked with 5% nonfat milk, probed with primary antibody, washed with TBST, followed by incubation with anti-rabbit HRP-conjugated secondary antibody. The signal was then detected by Clarity^TM^ western ECL substrate (Biorad).

### Metabolic Labeling and “Click” Chemistry

Cells were treated with 200 μM MalAM-yne or DMSO (vehicle control) for 1h. Cells were harvested, washed with PBS, and lysed. The lysate was incubated with the “click” reaction mixture (2 mM CuSO_4_, 3 mM THPTA, 40 mM ascorbate, and 2 mM TAMRA-azide) for 1 h at room temperature. The reaction was quenched with the addition of 4 volumes of ice-cold acetone. The reaction was kept at −20°C overnight, and centrifuged at 6,000 x *g* for 5 min at 4°C. The pellet was washed with ice-cold methanol twice, and air-dried for 10 min. The pellet was re-suspended in laemmli loading buffer. The samples were resolved by SDS-PAGE. To reduce the signal to noise ratio, the gel was destained in a mixture of methanol/distilled water/acetic acid (v/v/v = 4/5/1) to eliminate non-specific binding of free dyes. The destained gel and analyzed with in-gel fluorescence scanning using a Biorad ChemiDoc MP imager (excitation 532 nm, 580 nm cut-off filter and 30 nm band-pass). Commassie blue staining was applied to provide loading control.

### NR and NARH Treatment

Cells were incubated with NR or NARH at various concentrations for 6 h. Cells were harvested, rinsed and re-suspended in 1 mL of fresh medium. Cell number was determined using a hemocytometer. The cell suspension was then re-pelleted for NAD^+^ concentration determination.

### NAD^+^ concentration determination

To the cell pellet was added 30 μL of ice-cold 7% perchloric acid. The sample was then vortexed for 30 s and sonicated on ice for 5 min. The vortex-sonication cycle was repeated three times. The sample was centrifuged at 14,000 x *g* for 3 min at room temperature. Clear supernatant was removed and neutralized to pH 7 with 3 M NaOH and 1 M phosphate buffer (pH 9). The cellular NAD^+^ level was measured using NAD^+^ cycling assay as described previously.^42–44^

### Docking analysis

The crystal structure of SIRT5 complexed with NAD^+^ and H3K9Suc (PDB: 3RIY) was obtained from the RCSB Protein Data Bank.^5^ SYBYL-X 2.1 was used to prepare protein and small molecule for docking. The docking studies were performed using GOLD 5.6.8 program.^35^ NARH was docked in the binding sire, which was defined by the space in a 5 Å radius around R105. Docking conformations were scored by the default ChemPLP scoring function. The top solution was selected as the predicted binding pose.

## ACKNOWLEDGEMENTS

This work was supported in part by 1R01GM143176-01A1 from NIH/NIGMS (to Y.C.), VCU CCTR Endowment Fund (sub-award of UL1TR002649 from National Center for Advancing Translational Sciences to VCU) (to Y.C.), and the George and Lavinia Blick Research Fund (to Y.C.). MS analysis reported in this manuscript was performed at the Vermont Biomedical Research Network (VBRN) proteomics facility supported by P20GM103449 (NIH/NIGMS).

## Notes

### Competing Interest Statement

The authors have declared no competing interest.

