## Supplementary Figures for "Differential Regulation of SIRT5 Activity by Reduced Nicotinic Acid Riboside (NARH)"

**Synthesis of NARH**

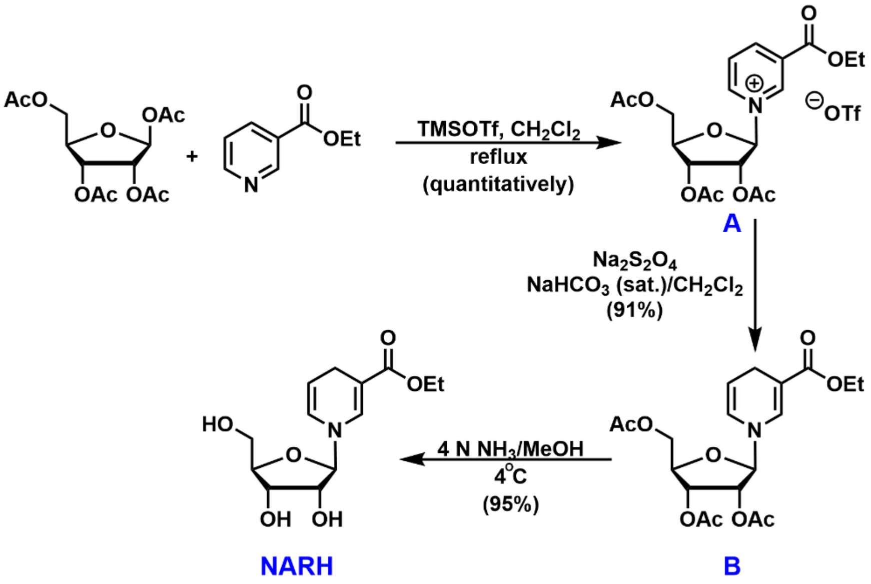


*Synthesis of Intermediate A*

Intermediate **A** was synthesized as described previously.[1, 2]

*Synthesis of Intermediate B*

In a round bottom flask flushed with argon, intermediate **A** (2.84 g, 5.08 mmol) was dissolved in 40 mL of $CH_2Cl_2$. To this solution was added 13 mL of saturated aqueous $NaHCO_3$ solution, followed by solid sodium dithionite (ca. 85%; 4.27 g; 20.9 mmol) and 7 mL of water. The biphasic reaction mixture was stirred at room temperature for 4 h, and then brine (30 mL) and $CH_2Cl_2$ (40 mL) were added. Yellow organic phase was separated, washed twice with brine, dried over $Na_2SO_4$, filtered and evaporated under reduced pressure to give a light yellow foam (intermediate **B**). The crude product was used for the next step without further purification.

*Synthesis of NARH*

Intermediate **B** (1.90 g, 4.62 mmol) was dissolved in 15 mL of methanol and 20 mL of 7N $NH_3$ in methanol at 0°C. The reaction was kept at 4°C for 16 h before solvent was removed under reduced pressure. The residue was dissolved in water and extracted with hexanes to remove organic impurities. The water layer was concentrated and purified by silica gel column chromatography ($CH_2Cl_2$ with 5%~10% methanol) to afford **NARH** (1.25 g, 4.37 mmol, 86% yield for two steps) as a pale yellow oil. $^1$H NMR ($CD_3OD$, 400 MHz) δ (ppm): 1.25 (t, *J* = 7.1 Hz, 3H), 3.04 (m, 2H), 3.61 (dd, *J* = 3.9, 12.0 Hz, 1H), 3.68 (dd, *J* = 3.4, 12.0 Hz, 1H), 3.84 (m, 1H), 4.00 (m, 2H), 4.12 (q, *J* = 7.1 Hz, 2H), 4.73 (d, *J* = 6.0 Hz, 1H), 6.10 (dd, *J* = 1.6, 8.2 Hz, 1H), 7.27 (s, 1H); $^{13}$C NMR ($CD_3OD$, 100 MHz): 14.89, 23.60, 61.02, 63.37, 72.16, 73.55, 85.81, 97.37, 99.69, 106.12, 126.64, 141.76, 170.43; HRMS (m/z): calculated for $C_{13}H_{19}NNaO_6$ (M+Na): 308.1105; found: 308.1114.

**Synthesis of MalAM-yne**

MalAM-yne was synthesized as reported previously.[3] $^1$H NMR ($CDCl_3$, 400 MHz) δ (ppm): 2.06 (t, *J* = 2.6 Hz, 1H), 2.14 (s, 6H), 2.84 (dd, *J* = 2.7, 7.6 Hz, 2H), 3.70 (t, *J* = 7.6 Hz, 1H), 5.80 (d, *J* = 1.7 Hz, 4H).

**Figure S1.** NMR and MS spectra of NARH.

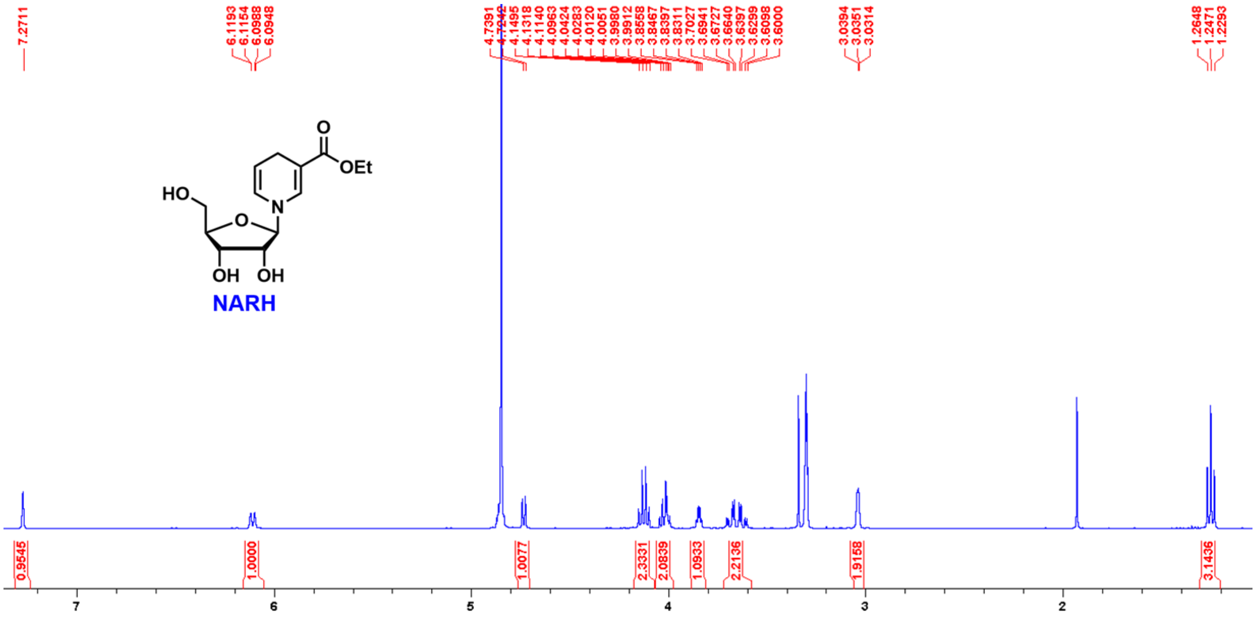


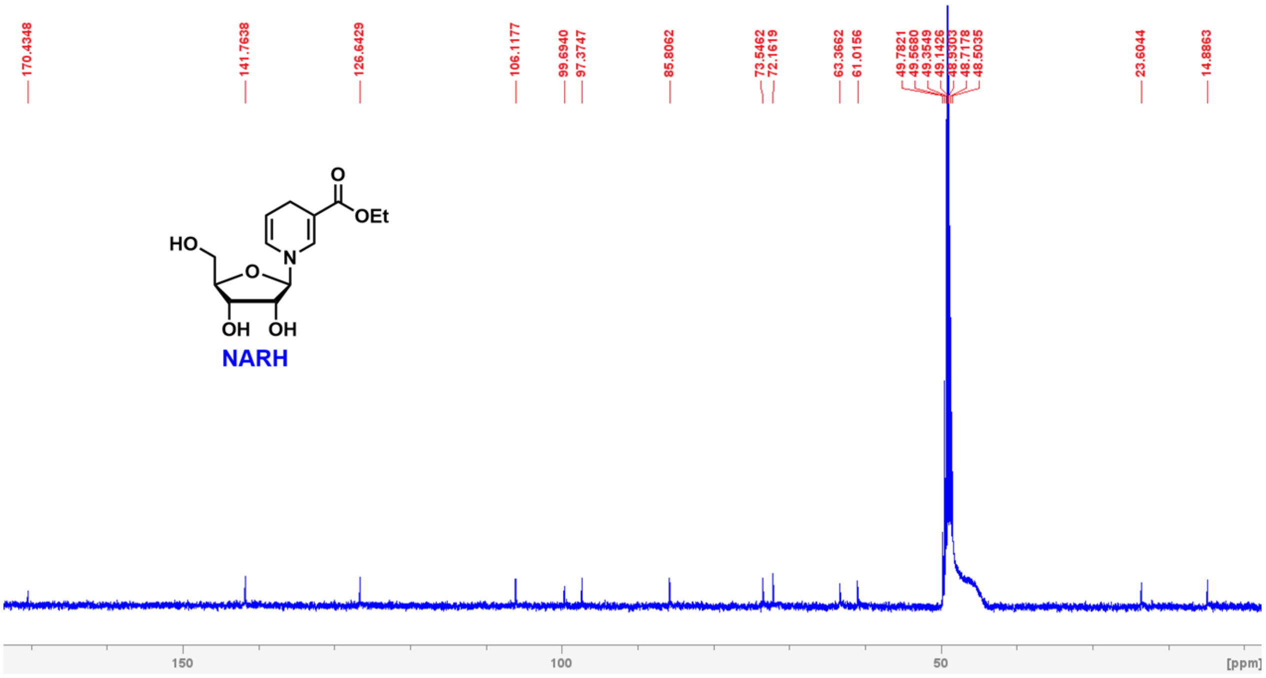

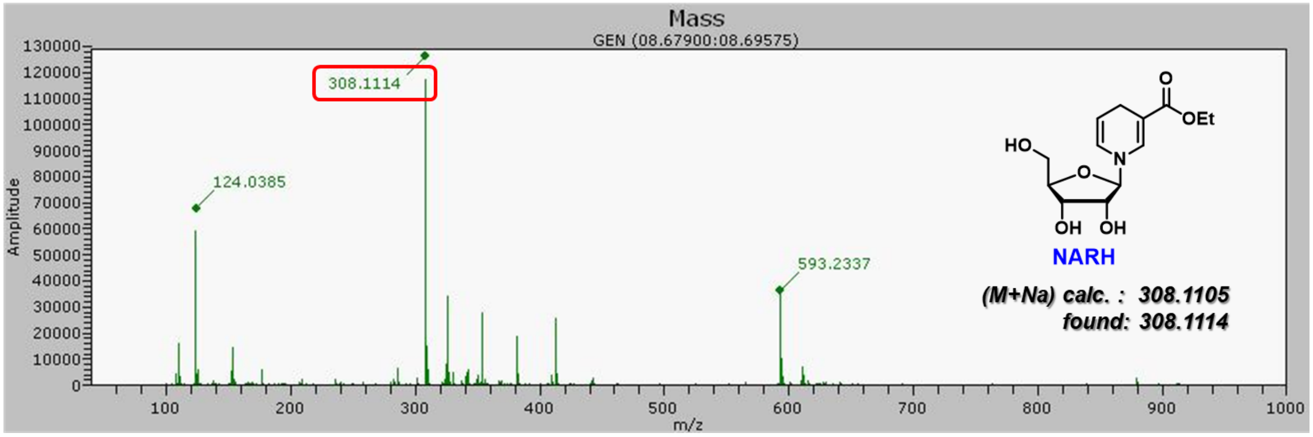

Mass
GEN (08.67900:08.69575)
Amplitude
m/z
308.1114
124.0385
593.2337
HO
O
OEt
N
OH
NARH
(M+Na) calc. : 308.1105
found: 308.1114

**Figure S2.** $^{1}$H NMR spectrum of MalAM-yne.

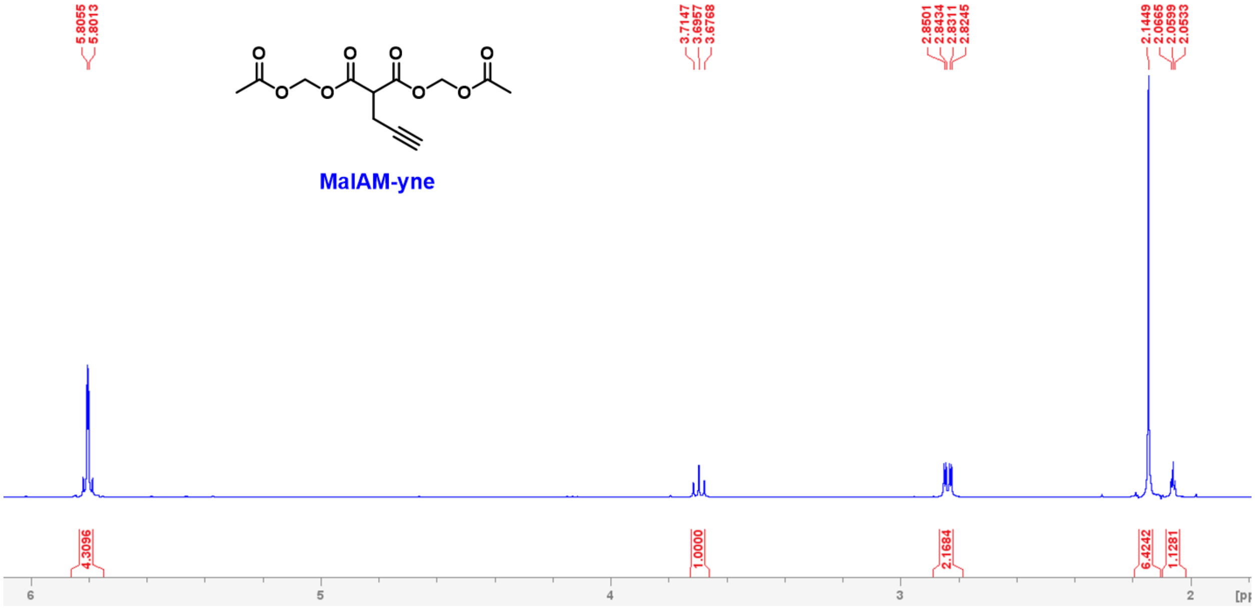

**Figure S3.** Dose-dependent activation of SIRT5 desuccinylation by NARH.

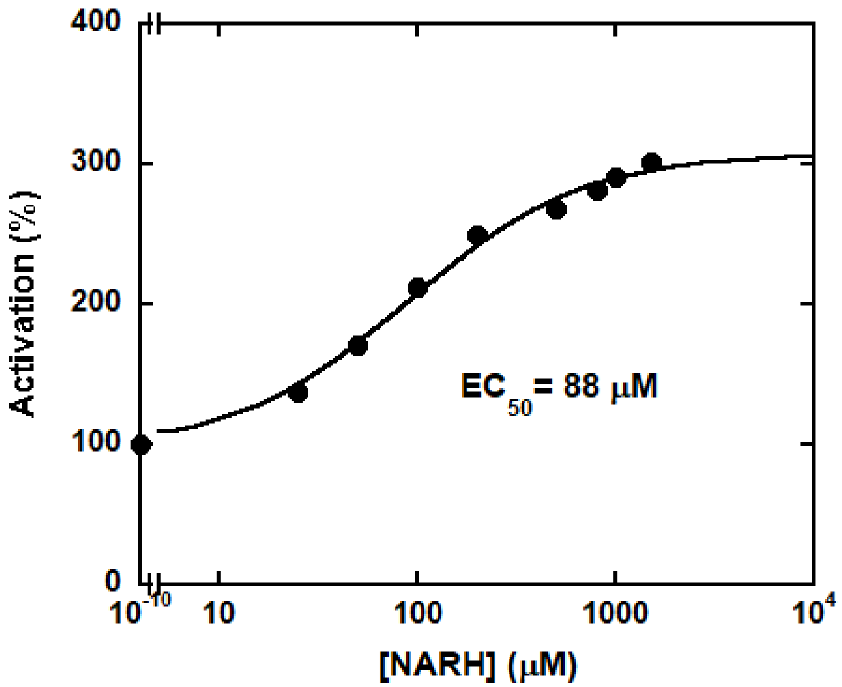


SIRT5 was incubated with $NAD^+$, H3K9Suc, and varying concentrations of NARH. The samples were then injected on an HPLC fitted to a Macherey-Nagel Nucleosil C18 column. Succinylated and desuccinylated peptides were resolved using a gradient of 10%–40% acetonitrile in 0.1% TFA. Chromatograms were analyzed at 215 nm. Reactions were quantified by integrating area of peaks corresponding to succinylated and desuccinylated peptides. The $EC_{50}$ was determined to be 88 μM.

**Figure S4.** NARH activates SIRT5 selectively.

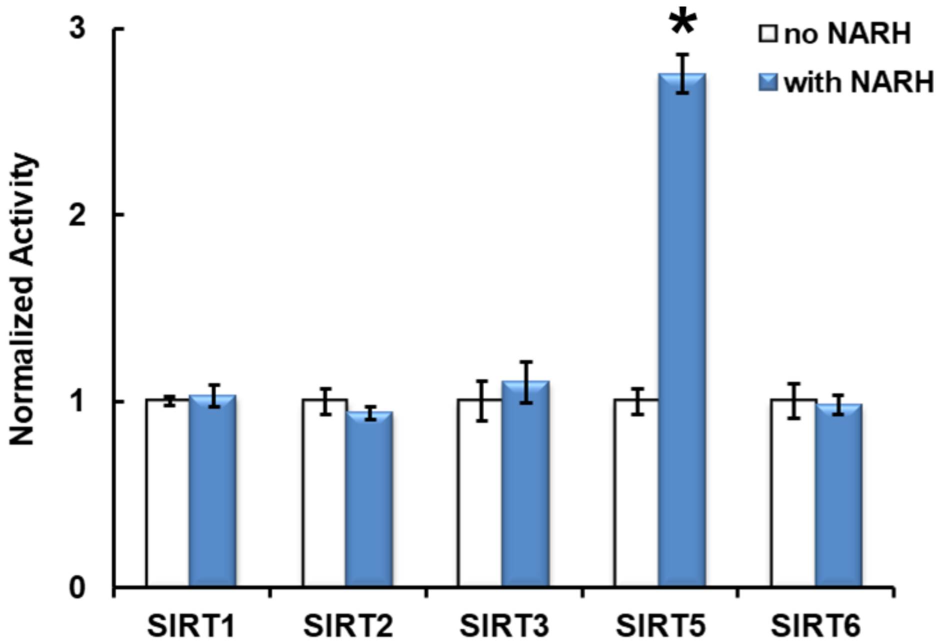


A typical reaction contained 800 μM $NAD^+$, 500 μM peptide substrate (H3K9Ac for SIRT2, SIRT3 and SIRT6, p53K382Ac for SIRT1, and H3K9Suc for SIRT5), with or without 800 μM NARH in 100 mM phosphate buffer pH 7.5. The reactions were initiated by the addition of 10 μM of sirtuin and were incubated at 37 °C before being quenched by 8 μL of 10% TFA. The incubation time was controlled so that the conversion of substrate was less than 15%. The samples were then injected on an HPLC fitted to a Macherey-Nagel Nucleosil C18 column. Acylated and deacylated peptides were resolved using a gradient of 10%–40% acetonitrile in 0.1% TFA. Chromatograms were analyzed at 215 nm. Reactions were quantified by integrating area of peaks corresponding to acylated and deacylated peptides. The data represents average of as least three independent experiments ± S.D. Statistical significance was determined by a Student's t-test: $*p < 0.01$.

**Figure S5.** ITC studies.

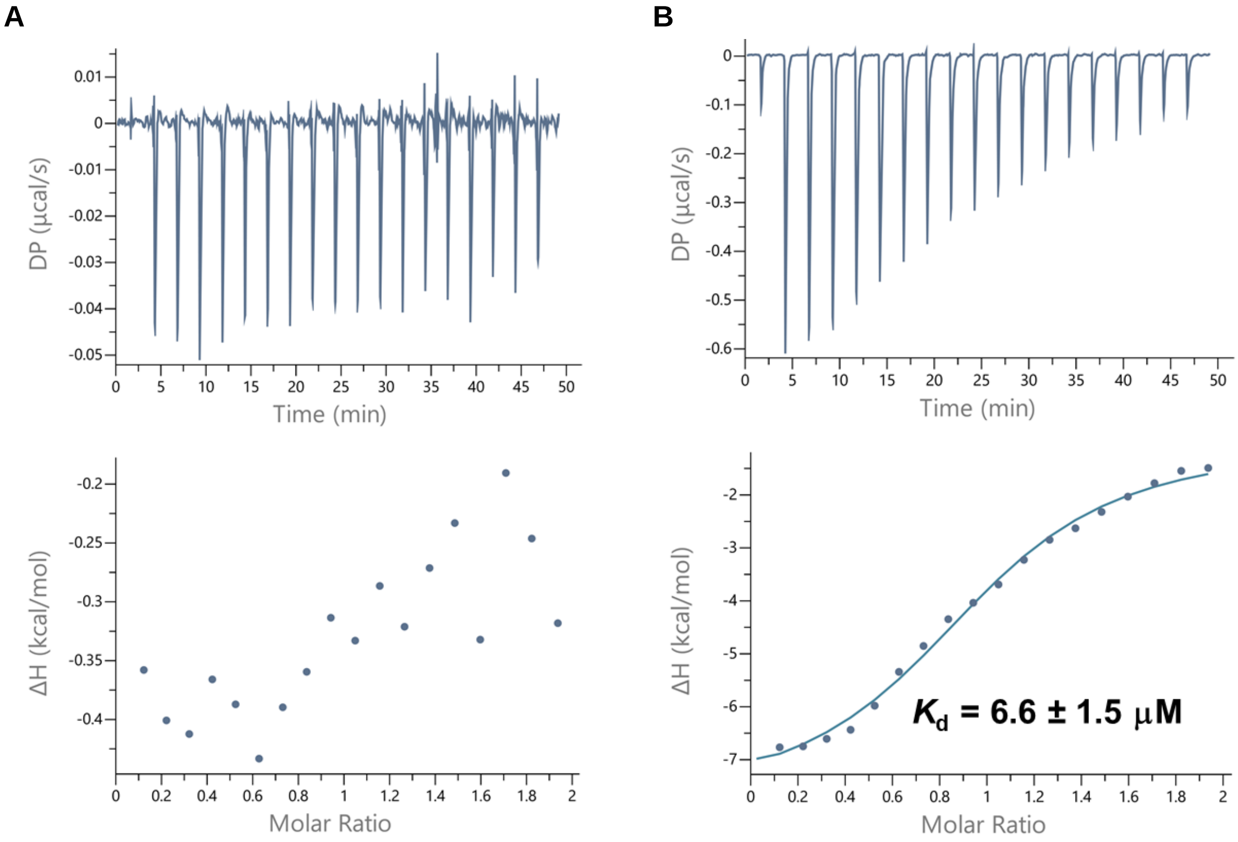


Representative ITC traces of $NAD^+$ (500 µM, A) or a mixture of $NAD^+$ and NARH (500 µM each, B) titrated into SIRT5 (50 µM). The experimental details can be found in "Methods and Materials".

**Figure S6.** NARH treatment does not increase cellular $NAD^+$ levels.

**A**

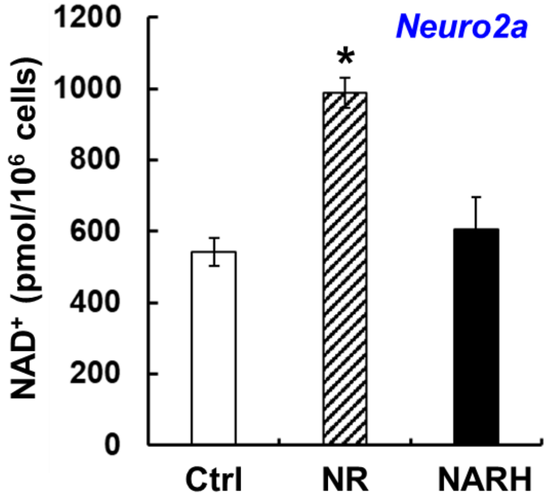


**B**

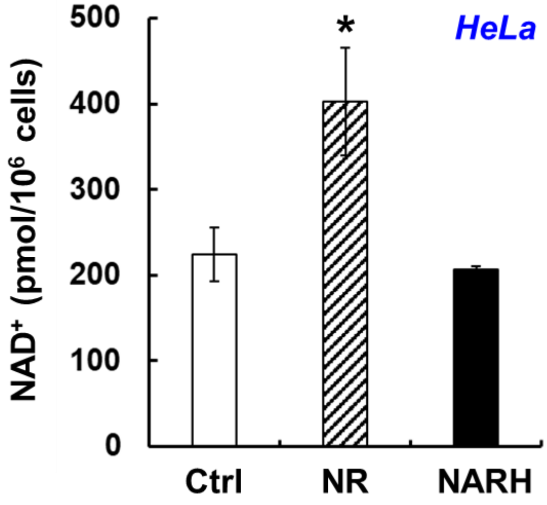


Intracellular $NAD^+$ levels in Neuro2a (A) and HeLa (B) cells. The cells were incubated with vehicle, 1 mM NR, or 1 mM NARH for 6 h. Subsequently, the intracellular $NAD^+$ concentrations were determined as described in “Methods and Materials”. The quantification data represents the average of three independent experiments ± SD. Statistical significance was determined by a Student’s *t*-test: *$p < 0.05$ *vs* control.

**Figure S7.** MST measurements of SIRT5 mutant binding to NARH.

A

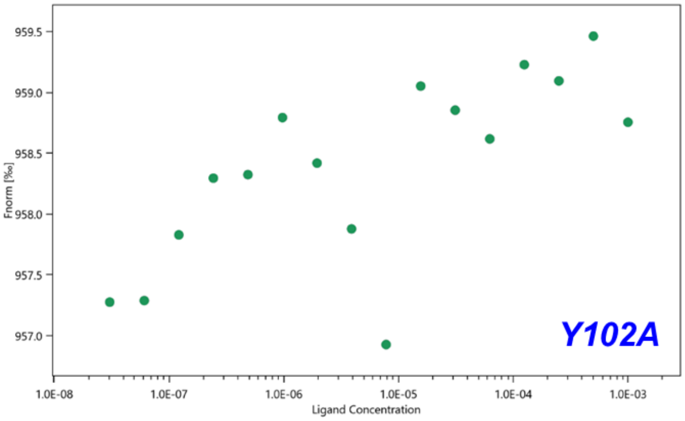


B

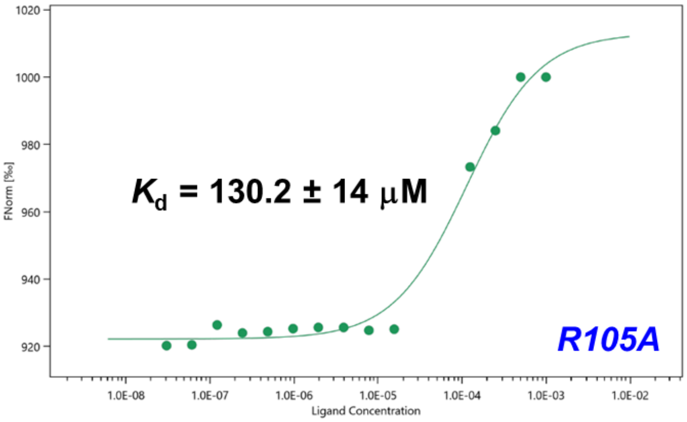


C

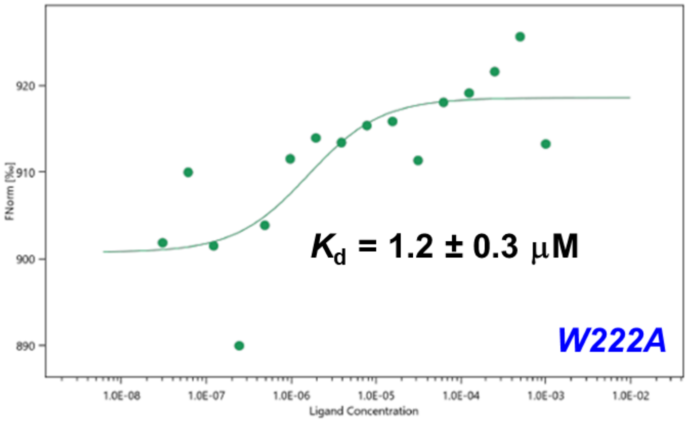

**Figure S8.** ADME prediction of NARH.

**Molecule 1**

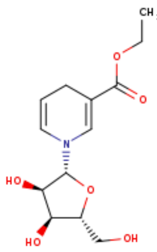


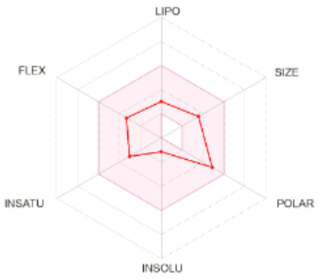


SMILES CCOC(=O)C1=CN(C=CC1)[C@@H]1O[C@@H]([C@H]([C@H]1O)O)CO

| Physicochemical Properties | |
|---|---|
| Formula | C13H19NO6 |
| Molecular weight | 285.29 g/mol |
| Num. heavy atoms | 20 |
| Num. arom. heavy atoms | 0 |
| Fraction Csp3 | 0.62 |
| Num. rotatable bonds | 5 |
| Num. H-bond acceptors | 6 |
| Num. H-bond donors | 3 |
| Molar Refractivity | 72.10 |
| TPSA | 99.46 Å² |
| **Lipophilicity** | |
| Log $P_{o/w}$ (iLOGP) | 2.45 |
| Log $P_{o/w}$ (XLOGP3) | -0.23 |
| Log $P_{o/w}$ (WLOGP) | -1.29 |
| Log $P_{o/w}$ (MLOGP) | -0.94 |
| Log $P_{o/w}$ (SILICOS-IT) | -1.14 |
| Consensus Log $P_{o/w}$ | -0.23 |

| Water Solubility | |
|---|---|
| Log *S* (ESOL) | -1.13 |
| Solubility | 2.10e+01 mg/ml ; 7.35e-02 mol/l |
| Class | Very soluble |
| Log *S* (Ali) | -1.40 |
| Solubility | 1.13e+01 mg/ml ; 3.97e-02 mol/l |
| Class | Very soluble |
| Log *S* (SILICOS-IT) | 1.03 |
| Solubility | 3.04e+03 mg/ml ; 1.07e+01 mol/l |
| Class | Soluble |
| **Pharmacokinetics** | |
| GI absorption | High |
| BBB permeant | No |
| P-gp substrate | No |
| CYP1A2 inhibitor | No |
| CYP2C19 inhibitor | No |
| CYP2C9 inhibitor | No |
| CYP2D6 inhibitor | No |
| CYP3A4 inhibitor | No |
| Log $K_p$ (skin permeation) | -8.20 cm/s |
| **Druglikeness** | |
| Lipinski | Yes; 0 violation |
| Ghose | No; 1 violation: WLOGP<-0.4 |
| Veber | Yes |
| Egan | Yes |
| Muegge | Yes |
| Bioavailability Score | 0.55 |
| **Medicinal Chemistry** | |
| PAINS | 0 alert |
| Brenk | 0 alert |
| Leadlikeness | Yes |
| Synthetic accessibility | 4.69 |

The physicochemical, ADME, and drug-likeness parameters of NARH were computed using the SwissADME online program (http://www.swissadme.ch/).[4]
